# Rescue of secretion of a rare-disease associated mis-folded mutant glycoprotein in *UGGT1* knock-out mammalian cells

**DOI:** 10.1101/2023.05.30.542711

**Authors:** Gábor Tax, Kevin P. Guay, Tatiana Soldà, Charlie J. Hitchman, Johan C. Hill, Snežana Vasiljević, Andrea Lia, Carlos P. Modenutti, Kees R. Straatman, Angelo Santino, Maurizio Molinari, Nicole Zitzmann, Daniel N. Hebert, Pietro Roversi, Marco Trerotola

**Affiliations:** Leicester Institute of Chemical and Structural Biology and Department of Molecular and Cell Biology, University of Leicester, Henry Wellcome Building, Lancaster Road, Leicester LE1 7HR, England, United Kingdom; Department of Biochemistry and Molecular Biology, and Program in Molecular and Cellular Biology, University of Massachusetts, Amherst, United States; Institute for Research in Biomedicine, Faculty of Biomedical Sciences, Università della Svizzera Italiana (USI), Bellinzona, Switzerland; Institute of Glycobiology, Department of Biochemistry, South Parks Road, Oxford OX1 3RQ, United Kingdom; Institute of Sciences of Food Production, ISPA-CNR Unit of Lecce, via Monteroni, I-73100 Lecce, Italy; Departamento de Química Biológica, Facultad de Ciencias Exactas y Naturales, Universidad de Buenos Aires (FCEyN-UBA) e Instituto de Química Biológica de la Facultad de Ciencias Exactas y Naturales (IQUIBICEN) CONICET, Pabellón 2 de Ciudad Universitaria, Ciudad de Buenos Aires C1428EHA, Argentina; Core Biotechnology Services, University of Leicester, University Road, Leicester LE1 7RH, England, United Kingdom; School of Life Sciences, École Polytechnique Fédérale de Lausanne, Lausanne, Switzerland; Institute of Agricultural Biology and Biotecnology, IBBA- CNR Unit of Milano, via Bassini 15, I-20133 Milano, Italy; Department of Medical, Oral and Biotechnological Sciences, “G. d’Annunzio” University of Chieti-Pescara, Italy; Laboratory of Cancer Pathology, Center for Advanced Studies and Technology (CAST), “G. d’Annunzio” University of Chieti-Pescara, Italy

**Keywords:** responsive mutant, glycoprotein secretion, UGGT, UGGT1, UGGT2, GDLD, Trop-2, *TACSTD2*.

## Abstract

Endoplasmic reticulum (ER) retention of mis-folded glycoproteins is mediated by the ER-localised eukaryotic glycoprotein secretion checkpoint, UDP-glucose glycoprotein glucosyl-transferase (UGGT). The enzyme recognises a mis-folded glycoprotein and flags it for ER retention by reglucosylating one of its N-linked glycans. In the background of a congenital mutation in a secreted glycoprotein gene, UGGT-mediated ER retention can cause rare disease even if the mutant glycoprotein retains activity (“responsive mutant”). Here, we investigated the subcellular localisation of the human Trop-2 Q118E variant, which causes gelatinous drop-like corneal dystrophy (GDLD). Compared with the wild type Trop-2, which is correctly localised at the plasma membrane, the Trop-2-Q118E variant is found to be heavily retained in the ER. Using Trop-2-Q118E, we tested UGGT modulation as a rescue-of-secretion therapeutic strategy for congenital rare disease caused by responsive mutations in genes encoding secreted glycoproteins. We investigated secretion of a EYFP-fusion of Trop-2-Q118E by confocal laser scanning microscopy. As a limiting case of UGGT inhibition, mammalian cells harbouring CRISPR/Cas9-mediated inhibition of the *UGGT1* and/or *UGGT2* gene expressions were used. The membrane localisation of the Trop-2-Q118E-EYFP mutant was successfully rescued in *UGGT1^-/-^*and *UGGT1/2^-/-^* cells. UGGT1 also efficiently reglucosylated Trop-2-Q118E-EYFP *in cellula*. The study supports the hypothesis that UGGT1 modulation constitutes a novel therapeutic strategy for the treatment of Trop-2-Q118E associated GDLD, and it encourages the testing of modulators of ER glycoprotein folding Quality Control (ERQC) as broad-spectrum rescue-of-secretion drugs in rare diseases caused by responsive secreted glycoprotein mutants.

**Synopsis:** Deletion of the *UGGT1* and *UGGT1/2* genes in HEK 293T cells rescues secretion of an EYFP-fusion of the human Trop-2-Q118E glycoprotein mutant. The mutant is retained in the secretory pathway in wild type cells and it localises to the cell membrane in *UGGT1*^-/-^ single and *UGGT1/2^-/-^* double knock-out cells. The Trop-2-Q118E glycoprotein disease mutant is efficiently glucosylated by UGGT1 in human cells demonstrating that it is a *bona fide* cellular UGGT1 substrate.

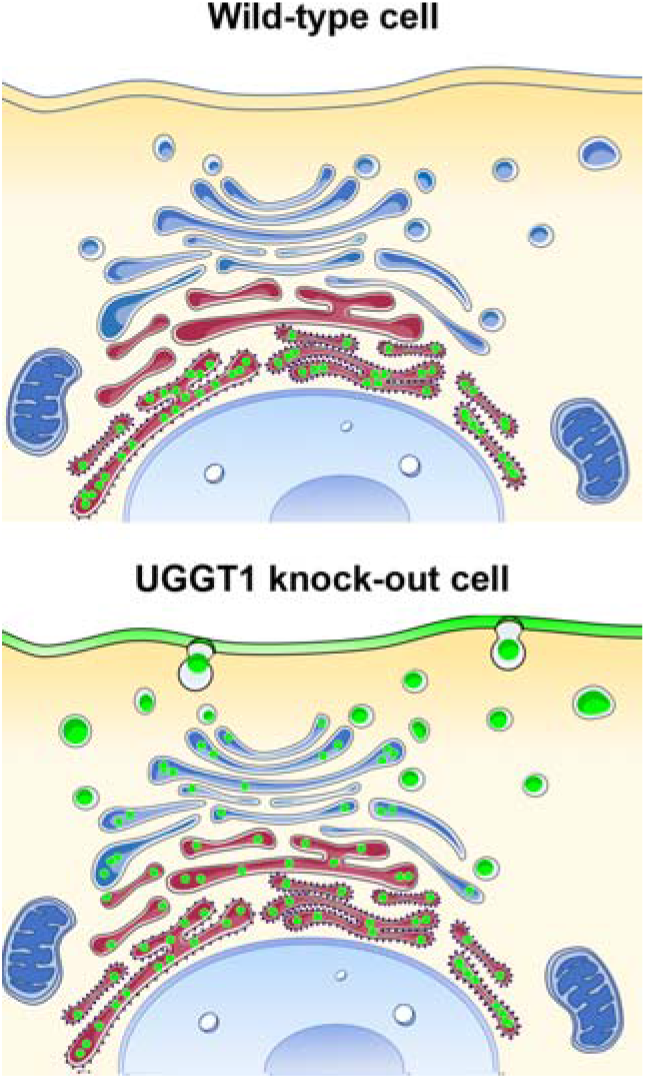

## Introduction

The Endoplasmic Reticulum (ER) glycoprotein folding Quality Control (ERQC) and Endoplasmic Reticulum Associated Degradation (ERAD) survey glycoprotein folding in the early secretory pathway. Both systems rely on checkpoint enzymes for detection of mis- folded glycoproteins. A glycoprotein that has not yet attained its native fold in the ER is recognised as mis-folded by the ERQC checkpoint enzyme, UDP-glucose glycoprotein glucosyltransferase (UGGT). UGGT flags a mis-folded glycoprotein for ER retention by reglucosylating one of its glycans and enables ER lectin-associated chaperones (calnexin and calreticulin) and foldases to aid its folding [1]. After prolonged cycles of UGGT-mediated ER retention, a terminally mis-folded glycoprotein in the ER is eventually recognised by the checkpoint of ERAD (a complex between an ER degradation-enhancing mannosidase, EDEM, and a protein disulfide isomerase, PDI), which de-mannosylates it, and dispatches it to retro-translocation and degradation in the cytosol [2]. Thus, ERQC/ERAD pathways provide means of coping with ER stress, ensure steady state glyco-proteostasis, prevent premature secretion of slowly folding glycoproteins and dispose of terminally mis-folded ones, thus conferring evolutionary advantages to healthy eukaryotic cells [3].

In the background of a secreted glycoprotein gene mutation that does not completely abrogate glycoprotein activity (“responsive mutant”, [4]), the stringency of the ERQC/ERAD machineries can be detrimental: UGGT recognises the mutated glycoprotein and flags it for ER retention, and the ERAD checkpoint de-mannosylates it, leading to degradation of the mutant glycoprotein, and to the concomitant loss of its residual activity. A variety of devastating rare diseases are indeed caused by overzealous ERQC/ERAD checkpoints preventing secretion of responsive mutant glycoproteins [5].

Replacement of the mutated gene with a wild type (WT) copy (gene therapy [6]) has the potential to help patients affected by congenital rare diseases. A second therapeutic strategy – whenever the patient’s mutation alters a gene encoding an enzyme, is enzyme replacement therapy [7]. A third strategy to help the subset of patients carrying responsive mutations is the use of a pharmacological chaperone [4]. A fourth, so far unexplored avenue for the therapy of congenital rare diseases caused by responsive glycoprotein mutants would be modulation of ERQC/ERAD checkpoint enzymes, which could act as rescue-of-secretion drugs [8,9]. Deletion of either the UGGT ERQC or ERAD misfolding checkpoints can rescue secretion of glycoprotein responsive mutants in the cress *Arabidopsis thaliana* [10,11]; inhibition of human ERAD mannosidases also rescues the secretion of a ß-sarcoglycan missense mutant [12]. No published study to date has investigated modulation of UGGT activity as a means of rescuing secretion of a human rare disease-associated responsive mutant glycoprotein.

Two paralogues of UGGT exist within the ER (UGGT1 and UGGT2). Only a few physiological clients of UGGT1 had been described [10,13–18] and the first partly overlapping subsets of UGGT1 and UGGT2 client glycoproteins (the “UGGT-omes” [19]) have only recently been characterised in HEK293 cells [20]. It is therefore not surprising that the role of UGGT in deciding the secretory trajectory of mis-folded glycoprotein mutants has been little studied in mammalian cells [9,14,21,22]. The mammalian UGGT1 isoform is essential during early embryogenesis and the homozygous *UGGT1*^-/-^ genotype causes mouse embryonic lethality, although cells derived from those embryos are viable [15]. Heterozygous *UGGT1*^+/-^ mice have been reported to express approximately half of the WT amount of UGGT1 and yet undergo normal development without any aberrant phenotype [23].

To date, there are no known UGGT selective inhibitors. UGGT is inhibited by its product UDP [24] and by squaryl derivatives of UDP [25]; by the non-hydrolysable UDP-Glucose (UDP-Glc) cofactor analogue UDP-2-deoxy-2-fluoro-D-glucose (U2F); and by synthetic analogues of the *N*-linked Man_9_GlcNAc_2_ glycan substrate [26,27]. Obviously, none of these molecules are UGGT-specific. A recently described UGGT1 and UGGT2 inhibitor was discovered in a fragment-based lead discovery effort [28] and is being chemically modified to increase its affinity and selectivity [29]. At the moment, *UGGT* gene knockout (KO), as a limiting case of inhibition, currently represents the only amenable strategy to probe the role of the enzyme in ER retention of glycoproteins.

In this manuscript, we describe rescue-of-secretion experiments of a congenital rare disease associated mutant of the membrane glycoprotein Trop-2 (*aka* epithelial glycoprotein- 1, gastrointestinal antigen 733-1, membrane component surface marker-1, or tumour-associated calcium signal transducer-2). This glycoprotein plays various roles in physiological and pathological contexts, among which cell-cell and cell-substrate interactions, cancer growth and metastasis [30–33]. Human Trop-2 (from now on indicated as Trop-2) is encoded by the *TACSTD2* gene, whose nonsense and missense mutations have been reported to cause gelatinous drop-like corneal dystrophy (GDLD, OMIM #204870) [34,35], a congenital rare disease that was first described in 1914 in a Japanese patient [36,37]. Since then, most of the GDLD cases were reported in individuals carrying mutations in the *TACSTD2* gene, in Japanese, Indian, Sudanese, Caucasian and Chinese populations, [38]. The worldwide prevalence of the disease is unknown; its frequency in Japan is estimated to be about 1/30,000, with about 4000 people affected by this disorder across the country: as is the case for the majority of congenital rare diseases, no cure is currently available for GDLD.

Patients with GDLD show a facilitated tear fluid permeation into their corneal tissue, and subepithelial amyloid deposition of lactoferrin; with ageing, the amyloid depositions typically enlarge, increase in number and coalesce with each other, leading to severe bilateral vision loss [39]. The *TACSTD2*^-/-^ mouse was recently developed and found to show a phenotype similar to that of human GDLD, although WT mice were also reported by the authors to develop corneal opacities [40]. Several missense mutations of the *TACSTD2* gene have been reported to cause GDLD [41], but given that only the *TACSTD2*^-/-^ mouse is available, alternative animal models are still required for fully investigated candidate responsive Trop-2 mutants *in vivo*.

Our experiments were conducted in WT and *UGGT1*^-/-^, *UGGT2*^-/-^ and *UGGT1/2^-/-^* HEK 293T and Vero E6 mammalian cells. Localisation of a fluorescent chimera of enhanced yellow fluorescent protein (EYFP) C-terminally fused to the WT and mutant Trop-2 was imaged by confocal laser scanning microscopy. For our *in cellula* study, we selected the GDLD-associated Trop-2-Q118E mutant [38] on the grounds that – upon modelling *in silico* the extracellular domain of Trop-2 and its missense mutants - the mutation seemed unlikely to impair function. In our experiments, we observe retention of Trop-2-Q118E-EYFP in WT and *UGGT2*^-/-^ cells, and rescue of secretion and plasma membrane localisation of the same mutant in *UGGT1*^-/-^ and *UGGT1/2^-/-^* cells. Pull-down experiments with recombinantly produced GST-tagged calreticulin (CRT) inhuman cell lysates using the KO background of dolichyl pyrophosphate Man_9_GlcNAc_2_ α-1,3-glucosyltransferase 6 (*ALG6^-/-^*) [20] demonstrated that the Trop-2-Q118E-EYFP mutant is indeed reglucosylated by UGGT1 *in cellula.* SDS-

PAGE and flow cytometry analysis of Trop-2-Q118E suggests that the majority of the secreted glycoprotein is aggregated via inter-molecular disulfides, making it unlikely that the mutant is responsive - as originally hypothesised. Yet, as it is likely that ERQC causes retention of many other rare-disease associated responsive glycoprotein mutants, our results encourage development and testing of modulators of ER glycoprotein folding as broad-spectrum rescue-of-secretion drugs in rare disease associated with responsive glycoprotein mutants.

## Results

### Homology modelling the Trop-2-Q118E mutant glycoprotein suggests mildly impaired protein folding

Distinct GDLD-causing missense mutations of Trop-2 have been reported. For our rescue of secretion study, we set out to select the least fold-disrupting of the known GDLD- causing missense mutations of Trop-2. In order to evaluate the likely impact of each mutation, we started by building a homology model of the dimer of the mature portion of human Trop-2 (Uniprot P09758, TACD2_HUMAN). Besides the `AlphaFold` Multimer model [42], a model was also built using Modeller [43]: the extracellular portion of the dimer (residues 27-268) was generated using as templates two distinct crystal structures of the cis-dimer of the same region of human Trop-2 (PDB IDs 7PEE [44] and 7E5N [45]); the transmembrane region (residues 278-298) was based on the NMR-derived model of the transmembrane portion of the rat p75 protein (PDB ID 4MZV, 30% sequence identity over 21 residues); the cytoplasmic domain of human Trop-2 (residues 299-323) was based on the NMR structure of the C-term of Trop-2 (PDB ID 2MAE). We then examined the likely consequences of GDLD-associated missense mutations by mapping them onto the Trop-2 homology model.

GDLD-associated mutations C108R and C119S are likely to cause severe misfolding accompanied by severe loss of function, as both Cys residues are involved in a disulfide bond in the extracellular domain. GDLD-associated mutations Y184C, L186P and V194E are also very likely to cause severe misfolding, as they will alter the protein hydrophobic core constituted by F180, F190, I203, L205, I219, A222 and F226. This area of the protein is distal to the membrane and it is involved in the trans dimer-of-dimers that establishes cell-cell adhesions [45]. The charge-reversal mutation E227K sits close to the dimer interface, surrounded by three positively charged residues: K72, K231 and R228: this is likely to destabilise the monomer and/or cause loss of protein dimerisation. The mutation Q118E appeared to be the best candidate for a putative responsive mutation: E is isosteric to Q, and the mutation affects only the surface area of the protein immediately above the membrane (see Figure S1).

### Molecular dynamics of WT and Q118E Trop-2 extracellular dimer suggests that the Q118E mutation results in destabilisation of tertiary and quaternary structure

In order to estimate the impact of the Q118E mutation on the Trop-2 dimer structure, we performed 1 µs all-atom Molecular Dynamics (MD) simulations of the extracellular domain of the WT Trop-2 dimer and of the corresponding Q118E mutant. The global stability of the extracellular domain dimers in time can be followed by measuring the root mean square deviation (rmsd_Cα_) of the protein backbone throughout the simulation, with the starting model taken as the reference. The Q118E mutant shows higher rmsd_Cα_ values than the WT, indicating that the mutation affects the stability of the protein fold (see Figure S2). In order to identify the regions most affected by the mutation, we measured the root mean square fluctuation (rmsf_Cα_) during the simulation, again using the starting structure as a reference. Both mutant and WT proteins show high values in N- and C-terminal regions. As expected, the thyroglobulin type-1 subdomain (TY, amino-acids 70 to 145) is the region most affected by the Q118E mutation (Figure S3). In the WT structure, the sidechain of Q118 forms several hydrogen bonds that stabilise the domain (Figure S1). The mutation introduces an extra negative charge that results in electrostatic repulsion between D107 and E118; the backbone O of R115 is also interacting with the side chain of (Q)E118. The distributions of the values of the distance between the alpha carbons of residues D107 and Q(E)118 during the simulation give an estimate of the destabilisation local to the TY subdomain introduced by the mutation (Figure S3).

### Secretion of the Trop-2-Q118E-EYFP mutant glycoprotein is impaired in WT HEK 293T and Vero E6 cells

In our *in silico* analysis, the Q118E mutation seemed the least likely to impair Trop-2 extracellular dimer structure in a severe manner. The mutant was therefore chosen as a candidate to test rescue-of-secretion of a rare-disease associated missense glycoprotein mutant by modulation of UGGT. In order to follow the subcellular localisation of the WT Trop- 2 glycoprotein and its Trop-2-Q118E mutant, WT and *UGGT* KO mammalian cells (HEK 293T, Vero E6) were transiently transfected with the Trop-2-pEYFP-N1 and Trop-2-Q118E- pEYFP-N1 plasmids, respectively encoding the EYFP C-terminal fusion of WT Trop-2 and its GDLD-associated Q118E mutant. Localisation of the fluorescent fusion glycoproteins was visualised 48 hrs post-transfection by confocal laser scanning microscopy imaging (Figures 1A, B). HEK 293T imaging used live cells, while Vero E6 imaging was carried out on fixed cells. Control experiments in cells transfected with a plasmid encoding Trop-2-pEYFP show the expected cell membrane localisation of the fusion glycoprotein, which traverses the secretory pathway and reaches the cell membrane both in WT and *UGGT1*^-/-^ cells (Figure 1A, white arrowheads). Conversely, in WT human HEK 293T and Vero E6 cells, the Trop-2- Q118E-EYFP mutant glycoprotein is retained in the secretory pathway (Figure 1B, right-hand side panels), a phenotype that mirrors Trop-2 lack of secretion in GDLD patients carrying the Trop-2 Q118E mutation.

**Figure 1.**
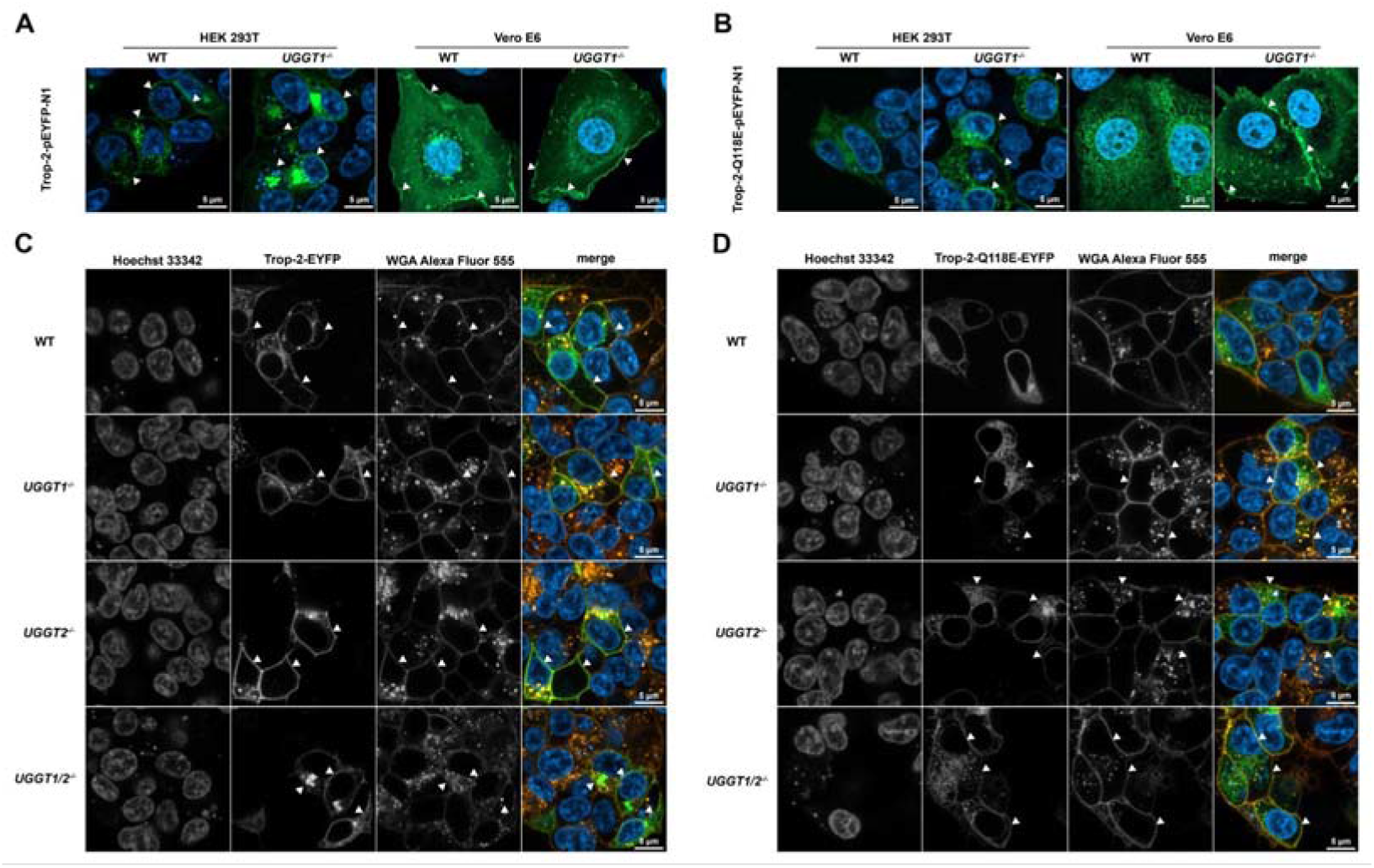
Rescue of secretion of the Trop-2-Q118E-EYFP- mutant glycoprotein upon UGGT1 inactivation. Confocal images of single optical sections of transiently transfected HEK 293T (live imaging) and Vero E6 cells (fixed cells). Nuclei were stained with Hoechst 33342 in HEK 293T cells or with DAPI stain in Vero E6 cells (blue). **(A, B)** Wild type (WT) and *UGGT1*^-/-^ HEK 293T and Vero E6 cells were transiently transfected with Trop-2-pEYFP-N1 and Trop-2-Q118E-pEYFP-N1 plasmids. **(A)** The Trop-2-EYFP wild-type glycoprotein (green) traverses the secretory pathway and reaches the plasma membrane (white arrowheads); **(B)** in WT cells, the Trop-2-Q118E-EYFP mutant glycoprotein (green) is visible in the secretory pathway but absent from the plasma membrane. In the *UGGT1*^-/-^ cells, the same Trop-2- Q118E-EYFP mutant glycoprotein reaches the cellular membrane (white arrowheads). **(C, D)** WT, *UGGT1*^-/-^, *UGGT2*^-/-^, and *UGGT1/2^-/-^* HEK 293T cells were transiently transfected with Trop-2-pEYFP-N1 **(C)** and Trop-2-Q118E-pEYFP-N1 **(D)** plasmids. Wheat Germ Agglutinin (WGA) Alexa Fluor 555 (orange) enables verification of plasma membrane localisation of the fluorescent fusion glycoproteins; **(C)** in both WT and *UGGT1^-^*^/-^ HEK 293T cells the Trop-2- EYFP-WT glycoprotein (green) reaches the cellular membrane (white arrowheads); **(D)** in the WT HEK 293T cells the Trop-2-Q118E-EYFP mutant glycoprotein (green) is trapped in the secretory pathway, while in the *UGGT1^-^*^/-^ and *UGGT1/2^-/-^*HEK 293T cells (and to a lower extent in *UGGT2*^-/-^ cells) the same mutant glycoprotein is visible both in the ER and in the membrane (white arrowheads).

### Secretion of the Trop-2-Q118E-EYFP mutant glycoprotein is rescued in *UGGT1* KO HEK 293T and Vero E6 cells

To assay membrane localisation of Trop-2 fluorescent glycoprotein, we stained cell membranes with wheat germ agglutinin (WGA) Alexa Fluor 555 conjugate (orange) (Figure 1C, D). Single KO (*UGGT1*^-/-^ and *UGGT2*^-/-^) and double KO (*UGGT1/2^-/-^*) HEK 293T cells were transiently transfected alongside WT HEK 293T control cells with Trop-2-pEYFP-N1 (Figure 1C, green) and Trop-2-Q118E-pEYFP-N1 plasmids (Figure 1D, green channel). The WGA fluorescence signal and the Trop-2-EYFP signal always overlap (Figure 1C), suggesting that Trop-2-EYFP reaches the membrane in both cell lines. The Trop-2-Q118E- EYFP mutant glycoprotein is absent from the membrane in the WT cell line (Figure 1D, top row). In the *UGGT1*^-/-^ and *UGGT1/2^-/-^* merged images, the green Trop-2-Q118E-EYFP reaches the membrane and overlaps with the WGA signal (orange, Figure 1D, white arrowheads): secretion of the Trop-2-Q118E-EYFP mutant glycoprotein is rescued in absence of UGGT1. Partial rescue of secretion of Trop-2-Q118E-EYFP is also detectable in *UGGT2*^-/-^ cells: the overlap of the green fluorescence with the Alexa Fluor 555 conjugate is less pronounced in *UGGT2*^-/-^ cells than in the *UGGT1*^-/-^ and *UGGT1/2^-/-^* cells (Figure 1D, white arrowheads).

### Multichannel intensity plot analysis confirms co-localisation of the Trop-2-Q118E-EYFP signal with the membrane dye in *UGGT1*^-/-^ KO HEK 293T cells

Multichannel intensity plots [46] were used to measure the overlap between the different fluorophores in the cellular membrane (Figure 2). We were able to validate our initial observations. The Trop-2-EYFP protein signal co-localises with the WGA Alexa Fluor 555 marker in the cellular membrane in all applied conditions (Figure 2, top panels). The Trop-2- Q118E-EYFP signal accumulates around the nucleus but not in the plasma membrane of WT HEK 293T cells (Figure 2, bottom left panel). In absence of *UGGT* (*UGGT1^-/-^*, *UGGT2^-/-^* and *UGGT1/2^-/-^*), the mutant glycoprotein localises at the plasma membrane (Figure 2, all bottom panels but the leftmost one), although to a lesser extent than in WT Trop-2 transfected cells.

**Figure 2.**
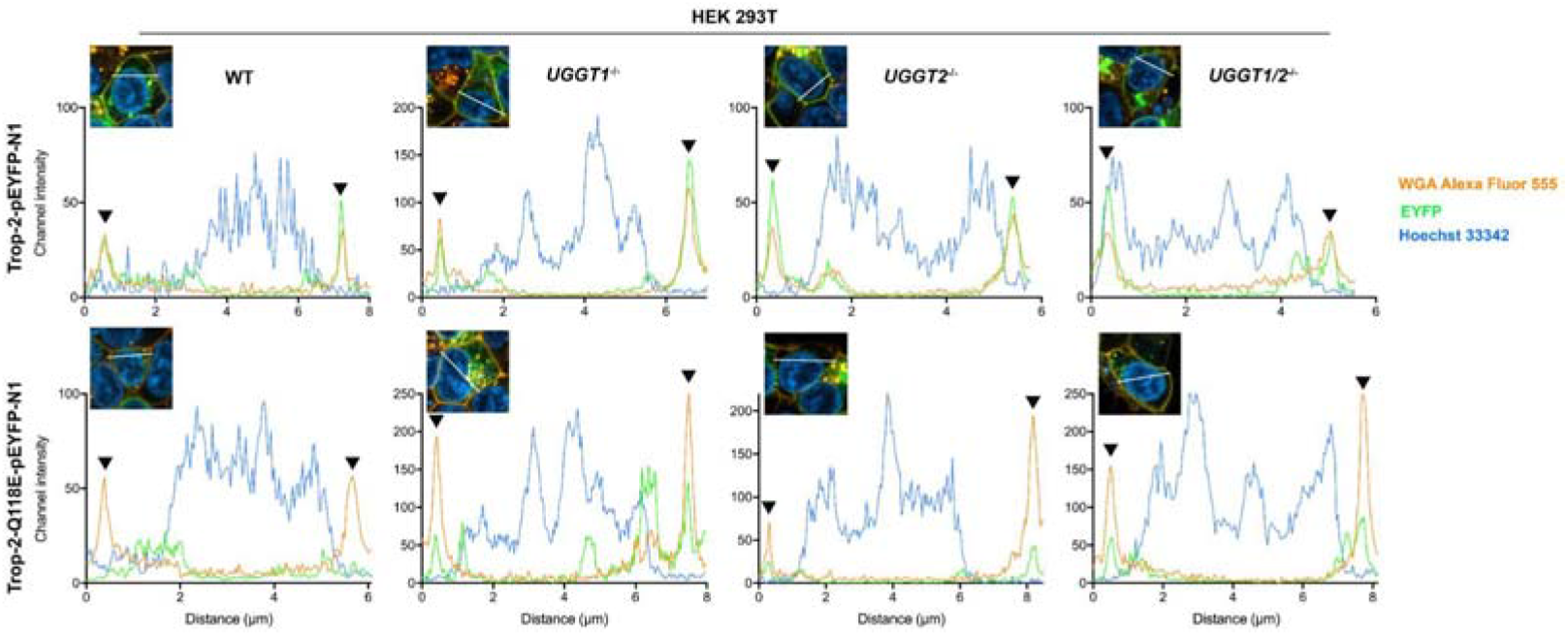
Multichannel intensity plots of the Trop-2-pEYFP-N1 transfected HEK 293T cell lines. For each combination of cell line and transiently transfected vector, the fluorescent signal is plotted along the line drawn in white in the inset panel. Labelling: green = EYFP; orange = WGA Alexa Fluor 555; blue = Hoechst 33342. The signal from the WGA Alexa Fluor 555 labelled membrane (orange) overlaps with EYFP (green) signals in Trop-2-pEYFP-N1 transfected cells (black arrowheads, top row). In case of the mutant Trop-2-Q118E-pEYFP- N1 transfected cells, the protein is absent from the membrane in WT cells (bottom left panel). The deletion of one or both *UGGT* genes rescues secretion of the mis-folded Trop-2 mutant glycoprotein (black arrowheads, all bottom panels but the leftmost one).

### The Trop-2-Q118E-EYFP lack of secretion phenotype is restored in HEK 293T *UGGT1*^-/-^ KO cells recomplemented with WT UGGT1 but not with inactive UGGT1

To check that the rescue of secretion observed for the Trop-2-Q118E-EYFP glycoprotein in the *UGGT1^-/-^* background was genuinely due to absence of UGGT1 catalytic activity in the KO cells, we performed UGGT1 re-complementation experiments. The previously generated WT and *UGGT1*^-/-^ KO HEK 293T cell lines were stably transfected with *Hs*UGGT1-pCDNA3.1/Zeo(+), encoding WT UGGT1, or with *Hs*UGGT1-D1454A- pCDNA3.1/Zeo(+) vectors, encoding an inactive mutant of UGGT1. The UGGT1 Asp1454 residue in this mutant is part of the conserved DQD motif in the catalytic domain [13,47] that likely bridges the substrate N-linked glycan and the UDP-glucose binding sites [13,29,48]).

The Trop-2-Q118E-EYFP mutant glycoprotein traverses the secretory pathway and reaches the cellular membrane only in UGGT1 D1454A re-complemented *UGGT1*^-/-^ cells (Figure 3B bottom row, and Figure 3C bottom row, third panel). In contrast to this, we observe the original lack of secretion phenotype and Trop-2-Q118E-EYFP glycoprotein retention in all cells expressing WT UGGT1 (Figure 3B, first three rows and Figure 3C bottom row, first, second and fourth panels). The localisation of the WT Trop-2-EYFP protein was not affected by the re-complementation (Figure 3A, and 3C top row panels). These results confirm that UGGT1 likely causes ER retention of the Trop-2-Q118E-EYFP mutant glycoprotein.

**Figure 3.**
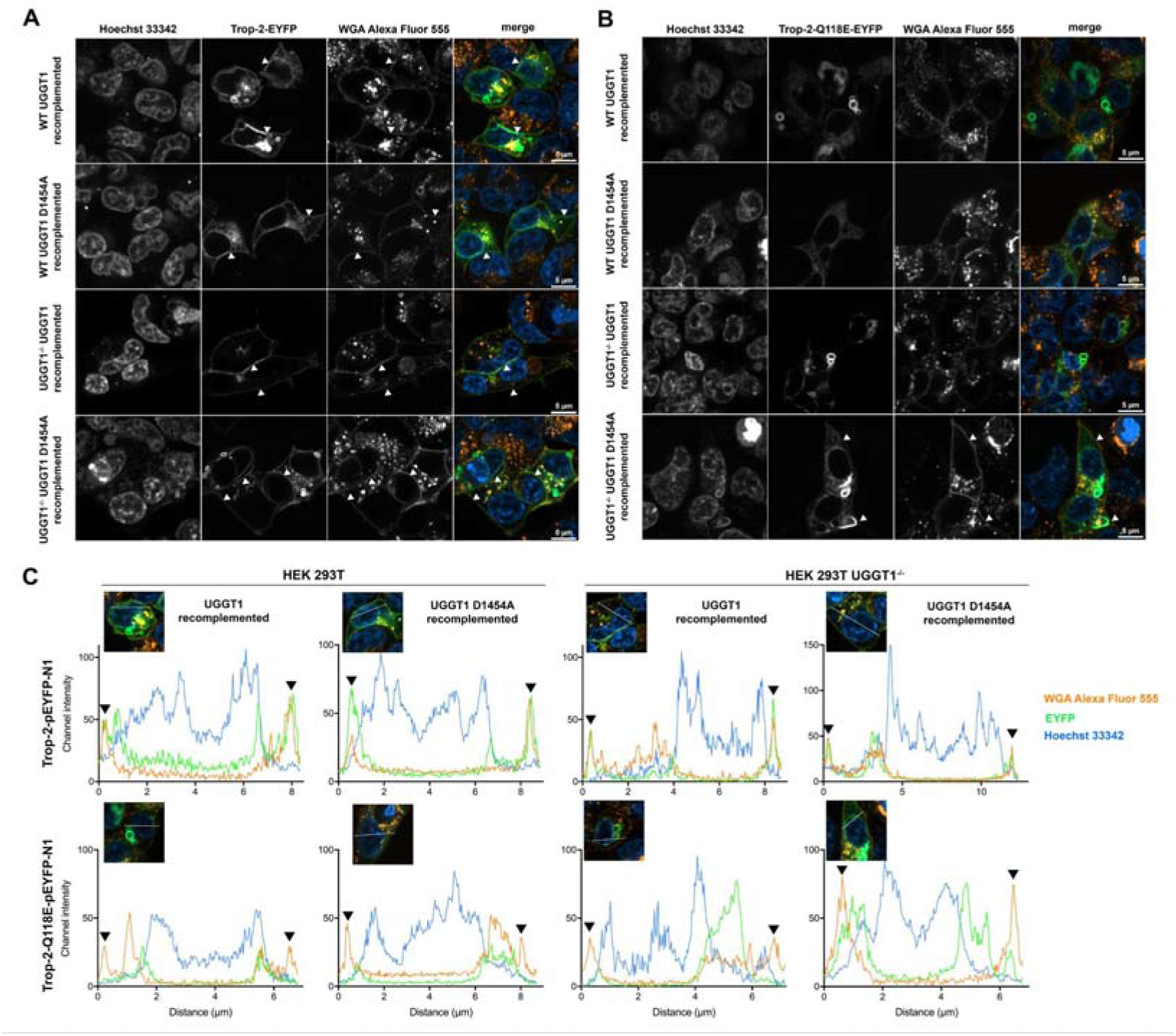
The Trop-2-Q118E-EYFP lack of secretion phenotype is restored in HEK 293T *UGGT1*^-/-^ cells re-complemented with the *UGGT1* gene. (**A**) UGGT1 and UGGT1 D1454A re-complementation does not interfere with the localisation of the WT Trop-2-EYFP glycoprotein (green): the protein traverses the secretory pathway and reaches the plasma membrane (white arrowheads); **(B)** In re-complemented WT cells, the Trop-2-Q118E-EYFP mutant glycoprotein (green) is absent from the plasma membrane and is retained in the secretory pathway (top first and second rows). In the UGGT1 re-complemented *UGGT1*^-/-^ cells, we observe restoration of the original lack of secretion phenotype observed for Trop-2- Q118E-EYFP in WT cells: the Trop-2-Q118E-EYFP mutant glycoprotein does not reach the cellular membrane (third row). Consistently with these observations, we still observe the rescue of secretion of the mutant glycoprotein in *UGGT1^-/-^* cells recomplemented with the inactive UGGT1 D1454A mutant (bottom row, white arrowheads). **(C)** Multichannel intensity plots of the Trop-2-EYFP transfected recomplemented cell lines confirm our visual observations (black arrowheads).

### The Trop-2-Q118E-EYFP mutant glycoprotein is efficiently glucosylated by UGGT1 *in cellula*

To confirm that ER retention of the mutant Trop-2-Q118E-EYFP glycoprotein is due to UGGT1, we assayed UGGT-mediated glucosylation of WT and mutant Trop-2-EYFP, transiently expressed into *ALG6^-/-^* HEK 293T cells. During the synthesis of *N*-linked glycans, as dolichol-phosphate linked precursors, ALG6 appends the first glucose to the immature Man_9_GlcNAc_2_ *N*-linked glycan precursor, which is further built up into the final tri-glucosylated form by additional ALG proteins within the ER (Figure 4A, [49]). In WT cells, calnexin and calreticulin (CRT) binding can occur either immediately after trimming of the Glc_3_Man_9_GlcNAc_2_ glycan to a Glc_1_Man_9_GlcNAc_2_ form, by glucosidases I and II, or through reglucosylation of a Man_9_GlcNAc_2_ glycan by one of the UGGTs [1]. Thus, any detected Glc_1_Man_9_GlcNAc_2_ glycans on glycoprotein clients in *ALG6^-/-^* HEK 293T cells must derive from UGGT glucosylation, and not from ER-glucosidase mediated partial *N*-linked glycan trimming (Figure 4B). The pool of UGGT-glucosylated glycoproteins was further enriched in our experiments in *ALG6^-/-^*HEK 293T cells by treating the cells with the glucosidase I and II inhibitor N-butyl-deoxynojirimycin (DNJ).

**Figure 4:**
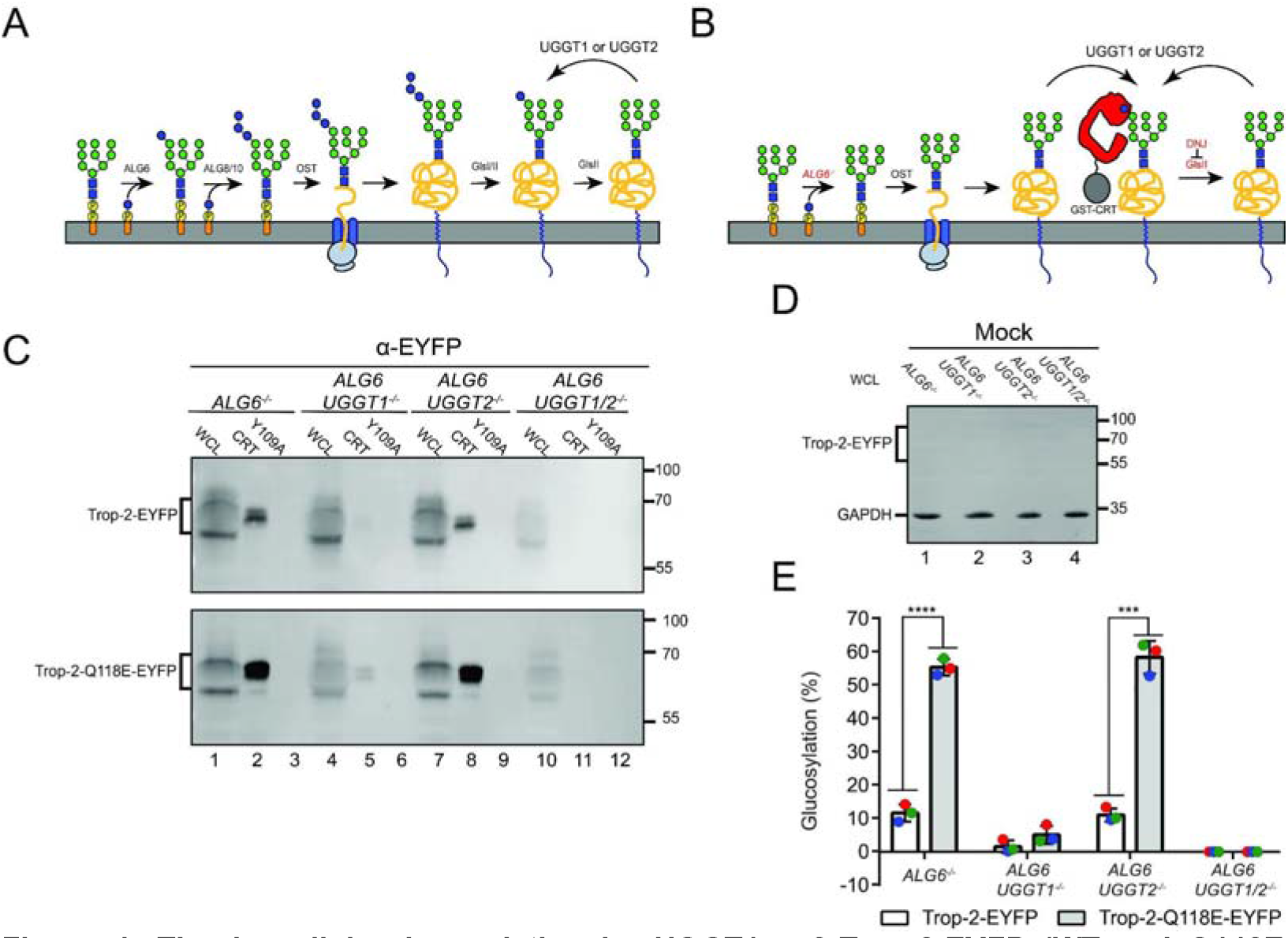
The *in cellula* glucosylation by UGGT1 of Trop-2-EYFP (WT and Q118E mutant). **(A)** The synthesis and modification of N-linked glycans starting from the dolichol-P- P-Man_9_GlcNAc_2_ stage within the ER lumen. ALG6 appends the first glucose onto the immature glycan using dolichol-P-glucose as the source. The mono-glucosylated glycan is then built into a tri-glucosylated carbohydrate by ALG8/10 prior to covalent linking to the nascent peptide chains. The glycan is trimmed to a mono-glucosylated state by glucosidases I and II, whereupon it can bind ER lectin chaperones. Removal of the remaining glucose by glucosidase II dissociates the glycoprotein from the lectin chaperones, whereby three options are possible: continuation of trafficking, degradation, or reglucosylation by the UGGTs. Orange ovals represent dolichol anchors, yellow circles phosphates, blue squares GlcNAc, green circles mannose, and blue circles glucose. **(B)** Alteration of glycan synthesis pathway for glucosylation studies. Deletion of *ALG6* results in a Man_9_GlcNAc_2_ glycan being transferred to nascent chains by the OST as the initial glucose can no longer be added. Interaction with ER lectin chaperones is thereby facilitated solely through glucosylation by the UGGTs. Further enrichment is achieved through treatment with DNJ, preventing the removal of the inner glucose. **(C)** The designated cell lines were transfected with either the WT Trop-2-EYFP or Q118E mutant and lysed. The lysate was split between a whole cell lysate sample (20%) and a GST-CRT (35%) and GST-CRT-Y109A pulldown (35%) and resolved by 9% SDS-PAGE before transferring to a PVDF membrane. Imaged is an αEYFP immunoblot. Data represent three independent biological replicates. **(D)** Mock transfection to measure background binding of αEYFP antibody against whole cell lysate from each cell line. The membrane was also probed against GAPDH as a loading control. **(E)** Quantification of blots from **(B)**. Percent glucosylation was calculated by subtracting the amount of protein pulled down in the Y109A lane from the CRT for each cell line. The resulting value was divided by the normalised quantified protein within the whole cell lysate and multiplied by 100. Error bars represent the standard deviation. **** represents P ≤ 0.0001 and *** represents P ≤ 0.001.

In addition to the single KO (*ALG6^-/-^)* cell line, previously generated double KOs (*ALG6/UGGT1^-/-^* and *ALG6/UGGT2^-/-^)* and triple KO (*ALG6/UGGT1/2^-/-^)* cells were also transiently transfected with Trop-2-pEYFP-N1 or Trop-2-Q118E-pEYFP-N1 vectors, to determine the levels of Trop-2 glucosylation and its specificity by either UGGT1 or UGGT2. Mono-glucosylated Trop-2 was isolated from the single, double and triple KO cell lines using a recombinant GST-tagged CRT that specifically binds mono-glucosylated glycoproteins. Any background or lectin-independent binding of mono-glucosylated glycoproteins was measured and subtracted using a lectin-deficient variant of CRT, CRT(Y109A) [50]. The adjusted value was divided by the amount of substrate within the whole cell lysate (WCL) quantifying the percent glucosylation (Figure 4C and E, [20]).

Both the WT and Q118E mutant Trop-2 are glucosylated by UGGT (Figure 4C, lane 2) and are substrates specifically of UGGT1 (Figure 4C, lane 8). Minimal glucosylation is attributed to UGGT2 (Figure 4C, lane 5). As expected, no glucosylation is observed in the triple KO (Figure 4C, lane 12). Of note, no lectin-independent interactions are observed in any CRT(Y109A) control pulldowns (Figure 4C, lanes 3, 6, 9, 12).

The Q118E substitution significantly increases the recognition and glucosylation by UGGT1 in both *ALG6^-/-^* and *ALG6/UGGT2^-/-^*cell lines (Figure 4C, lanes 2, 8 and Figure 4E). Levels of glucosylation increase from ∼10% to ∼60%, for the WT and Q118E mutant respectively, in both the *ALG6^-/-^* and *ALG6/UGGT2^-/-^* cells. Minimal glucosylation is observed within the *ALG6/UGGT1^-/-^*cell line with no significant difference being measured. Taken together, these results demonstrate that both the WT and mutant Trop-2 are efficiently glucosylated within the ER and the Q118E mutation significantly increases the amount of glucosylation by UGGT1. This suggests that the likely mechanism for the failure of the mutant to traffic to the cell surface is through increased reglucosylation by UGGT1, and subsequent lectin chaperone binding, leading to ER retention of the mutant.

### The Q118E mutation results in increased inter-molecularly SS-bonded mis-folded Trop- 2 multimers

To acquire insight on the nature of the misfolding recognised by UGGT1 in the Trop- 2-Q118E-EYFP mutant, we analysed (by Western blotting with an anti-EYFP antibody) membrane fractions from cells transiently transfected with the Trop-2 WT and Q118E mutant glycoproteins. Membrane fractions were isolated with styrene maleic acid lipid particles (SMALPs, Lipodisq, Figure 5A) or with CelLytic MEM Protein Extraction Kit ™ (Figure 5B). Inter-molecularly SS-bonded Trop-2 multimers are detected in small amounts in all membrane samples under non-reducing conditions. The amounts of these inter-molecularly SS-bonded multimers increase when the cells express the Trop-2-Q118E mutant (Figure 5B, left-hand side).

**Figure 5.**
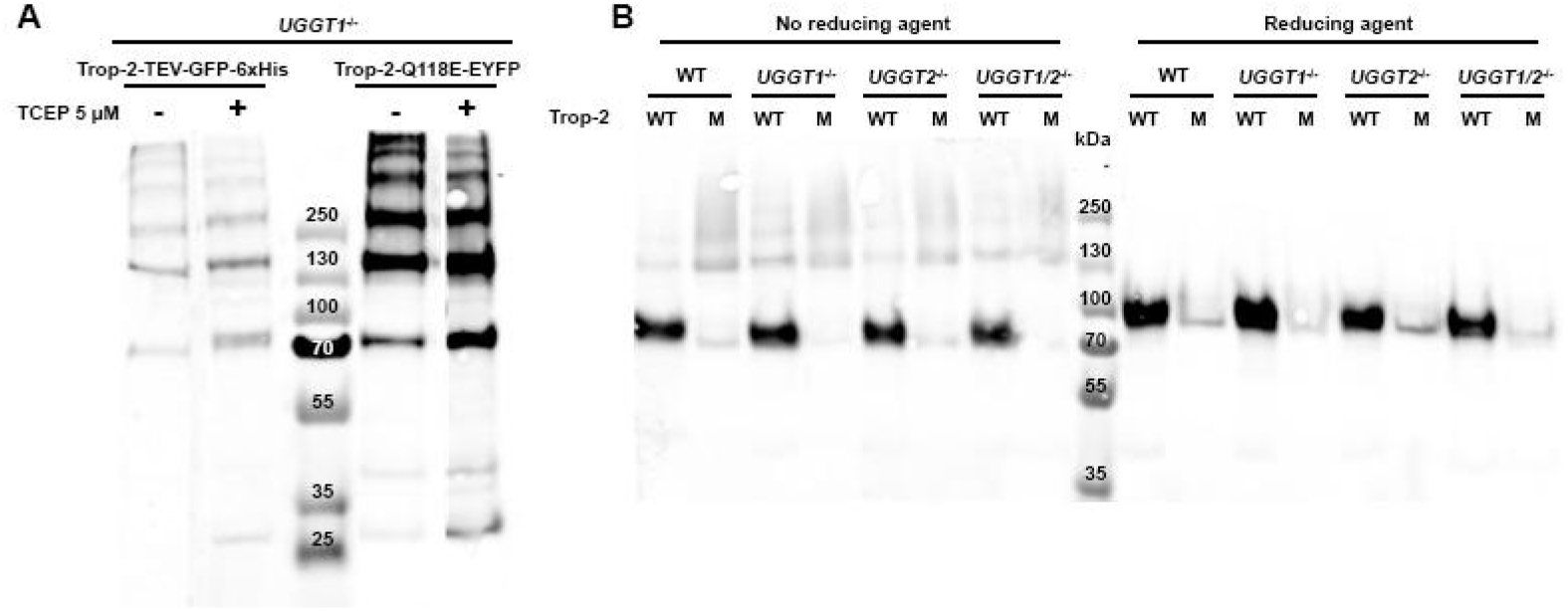
The Q118E mutation generates inter-molecularly SS-bonded mis-folded Trop- 2-Q118E-pEYFP-N1 multimers. **(A))** Anti-EYFP Western blot of membrane proteins isolated by Lipodisq extraction 96 hrs post-transfection from HEK 293T *UGGT1^-/-^*cells transiently transfected with Trop-2-TEV-GFP-6xHis WT and the Trop-2-Q118E mutant in the presence or absence of TCEP. **(B)** Anti-EYFP Western blot of membrane fractions extracted with the CelLytic™ MEM Protein Extraction Kit from HEK 293T cells transiently transfected with the Trop-2-pEYFP-N1 (“WT”) and Trop-2-Q118E-pEYFP-N1 plasmids (“M”). Samples were run in the abeance or presence of reducing agent..

### Multiparametric flow cytometry confirms that secretion of mis-folded Trop-2-Q118E mutant is rescued in *UGGT1^-/-^*cells

Biochemical analysis suggested that the majority of the secreted Trop-2-Q118E mutant is aggregated via inter-molecular disulfides. We further probed the structural integrity of the rescued Trop-2 mutants by multiparametric flow cytometry on *UGGT1^-/-^* Vero E6 cells expressing WT or Q118E Trop-2-EYFP fusion glycoproteins. Specifically, we stained these cells with two different types of far-red (APC) labelled antibodies: a conformation-insensitive goat polyclonal antibody (pAb, AF650 from R&D) and the T16 mouse monoclonal antibody (mAb), which only binds the native/correctly folded Trop-2 [51]. We gated out EYFP^+^ cells, and compared the mean fluorescence intensities (MFIs) of the cells stained with mAb and pAb. As shown in Figure 6, the mAb/pAb MFI ratio was 1.63 in the cells expressing Trop-2- EYFP, while it was 0.21 in the cells expressing Trop-2-Q118E-EYFP. This analysis confirms that the secretion of the Trop-2-Q118E mutant is rescued upon KO of *UGGT1*, although the mutant is to some extent structurally compromised.

**Figure 6.**
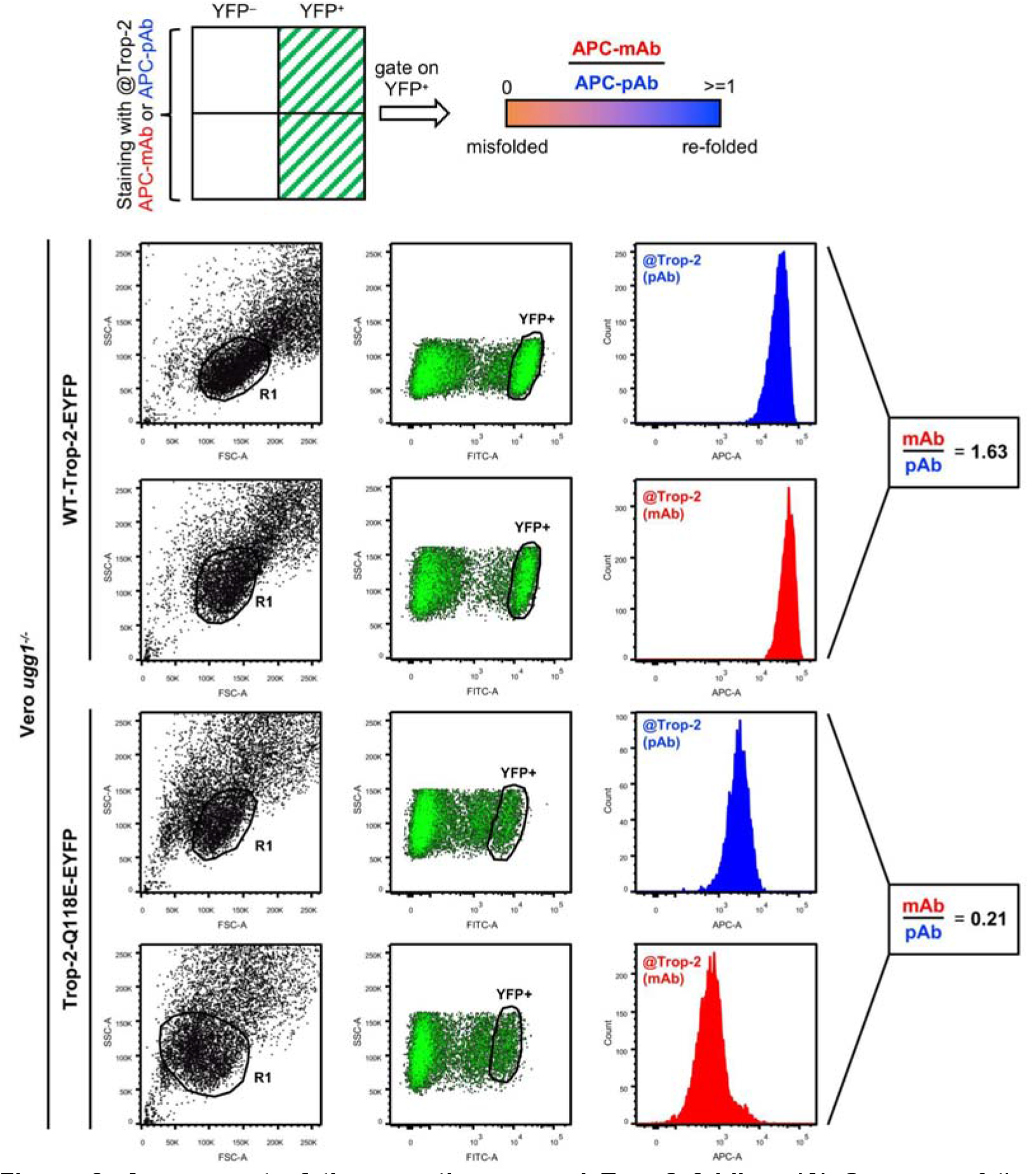
Assessment of the secretion-rescued Trop-2 folding. **(A)** Summary of the gating and signal acquisition strategy. **(B)** Left panels: dot plots of the experiments showing dimensional parameters (forward scatter, FSC, and side scatter, SSC). *UGGT1^-/-^* Vero E6 cell singlets (R1) were gated out for subsequent analysis of *UGGT1^-/-^* Vero E6 cells expressing WT or Q118E-Trop-2-EYFP proteins. Middle panels: Fluorescent signals in the FITC/YFP channel relative to the R1 cells are shown. YFP^+^ cells were gated out for following analysis. Right panels: histogram plots showing the expression levels of membranous Trop-2 as measured by staining with an anti Trop-2 pAb (blue) or mAb (red). Ratios between mAb and pAb Mean Fluorescence Intensities are shown for each experimental group.

## Discussion

In the background of a congenital mutation in a gene encoding a secreted glycoprotein, the ER glycoprotein misfolding checkpoint enzymes (UGGTs and EDEMs) are likely responsible for prolonged ER retention and eventual degradation of the mutant glycoprotein [52]. This in turn gives rise to loss-of-function congenital rare disease. Thus, for a wide range of slightly mis-folded but still functional glycoproteins (“responsive mutants” [4]), the fidelity of ERQC/ERAD-mediated quality control paradoxically exacerbates the severity of disease [5,12,53]. Evidence of disease-associated lack of secretion exists for several rare disease associated glycoprotein mutants (*e.g.* [54–57]). For patients carrying glycoprotein mutants that fail to reach the intended cellular or extracellular destination, but retain measurable residual activity, enzyme replacement [7] and/or pharmacological chaperone therapy [58] are viable options. Unfortunately, databases of rare disease associated congenital missense mutations do not routinely report residual activity information. Thus, the scope of rescue-of-secretion therapy - in terms of overall number of patients carrying responsive mutations across the full spectrum of congenital rare disease - is difficult to assess.

For the subset of congenital rare disease patients carrying a responsive mutation, modulation of the ER glycoprotein misfolding checkpoint enzymes (UGGT/EDEM) is an alternative, yet-to-be-explored strategy to restore mis-folded active glycoprotein secretion and alleviate the disease symptoms [8,9,12]. For all glycoprotein clients that are either slow to fold and/or carry toxicity when mis-folded, modulation of UGGT/EDEM of course carries an intrinsic risk of premature and unwanted secretion, a scenario we dubbed the “ER Pandora’s box” [19]. Furthermore, no specific inhibitors exist for either UGGTs or EDEMs, so that the full impact of UGGT/EDEM modulation on the (glyco)secretome of cells with pharmacologically loosened ERQC/ERAD is still difficult to assess. Perhaps for these reasons, rescue of secretion of mis-folded glycoprotein mutants by pharmacological UGGT/EDEM modulation has not been pursued and the molecular basis for the lack of secretion of glycoprotein mutants has been investigated only in a few cases [9,15,21,22].

Recently, the first set of UGGT1 and UGGT2 client glycoproteins in HEK293 cells (the HEK293 “UGGT-omes” [19]) have been discovered [20], and a sub-millimolar inhibitor of both isoforms described [29], paving the way for investigation of the cellular consequences of UGGT modulation. The only rescue-of-secretion experiments of responsive glycoprotein mutants by UGGT KO published so far are the ones in plants, but ERQC/ERAD are well conserved in eukaryotes. We decided to test rescue of secretion in UGGT knock-out mammalian cells by monitoring the localisation of a WT and mis-folded rare-disease associated glycoprotein mutant by confocal microscopy, testing *UGGT1*^-/-^, *UGGT2*^-/-^ and *UGGT1/2^-/-^* HEK 293T and Vero E6 cell lines.

For the rescue-of-secretion experiments, amongst the many candidates for rare-disease associated mis-folded glycoprotein mutants, we choose to follow fluorescently labelled human Trop-2 glycoproteins. Membrane localisation of WT Trop-2 does not appear to depend on the presence of either UGGT isoforms: the WT glycoprotein reaches the cell membrane in all cell lines tested, with no statistical differences observed in the degree of co- localization with a cell membrane marker (Figures 1 and 2).

In contrast to WT Trop-2, the Trop-2-Q118E mutant glycoprotein does not reach the cell membrane in WT HEK 293T and Vero E6 cells (Figure 1). Lack of secretion likely underpins the GDLD associated with the mutation. Rescue of secretion of the same mutant in the *UGGT1*^-/-^ and *UGGT1/2^-/-^*cell lines (Figures 1 and 2) suggests a role for UGGT1 in recognising a folding defect. Re-complementation of the *UGGT1*^-/-^ and *UGGT1/2^-/-^* cell lines with a WT *UGGT1* gene reverts the phenotype to the lack of secretion observed in the WT cells, while recomplementation of the same with an inactive *UGGT1* mutant still shows rescue of secretion of the mis-folded Trop-2 mutant (Figure 3). These data support the hypothesis that UGGT1 activity is the source of ER retention of the mutant seen in the WT cells.

In order to confirm that the rescue of secretion observed in the *UGGT1*^-/-^ background is due to Trop-2-Q118E mutant glycoprotein being a client of UGGT1 (and not to an indirect effect of the *UGGT1* KO) we went on to test *in cellula* reglucosylation of Trop-2-Q118E. Our results confirm that Trop-2-Q118E is indeed a UGGT1 client (Figure 4). In our experiments, retention of the Trop-2-Q118E glycoprotein mutant in *UGGT2*^-/-^ cells appears statistically equivalent to the WT one, but rescue of secretion in the *UGGT1/2^-/-^* cells is slightly improved over the one observed in the *UGGT1*^-/-^ background. These data may suggest a role for UGGT2 in the recognition of the defect in Trop-2-Q118E and/or in the folding of glycoproteins that mediate ER retention of the mutant. Indeed, glucosylation of the Trop-2-Q118E glycoprotein mutant by UGGT2 is also detectable *in cellula* (Figure 4), albeit at much reduced levels when compared with UGGT1-mediated glucosylation.

Despite the molecular determinants underpinning misfold-recognition by UGGT remaining elusive yet, it is currently hypothesised that the enzyme recognizes and binds to mis-folded glycoproteins through specific structural features, such as hydrophobic patches exposed on the protein surface due to incorrect folding. Using MD simulations we found that Q118E mutation induces a disruption of the Trop-2 fold, likely due to an electrostatic repulsion between D107 and the sidechain of E118. This would result in a loss of the overall stability of the protein and transform the mutant glycoprotein into a substrate for UGGT.

Our Western blots detect intermolecular SS-bonded Trop-2 multimers when overexpressing the protein in HEK293 cells (Figure 5). The amount of Cys mispairing is significantly increased by the Q118E mutation. Membrane inserted Trop-2 is a dimer in which each monomer has five intramolecular disulphide bridges. In particular, Q118 is flanked by the C119-C125 disulfide bridge (see Figure S1), and the Q118E mutation may disturb its formation. Consistent with this evidence, our flow cytometry experiments carried out using anti Trop-2 antibodies show that a large fraction of the Q118E mutant whose secretion is rescued in *UGGT^-/-^*cells does not fold properly.

The fraction of rare disease patients carrying a mild missense mutation in a secreted glycoprotein gene is not known, and it varies for each glycoprotein/rare disease. For example, in lysosomal storage diseases and cystic fibrosis, an estimated 15-20% and 70% of patients carry a responsive mutant [4]. The overall scope of ERQC/ERAD modulation for the therapy of such patients is yet to be assessed. Each of these patients’ mutation is in principle amenable to pharmacological chaperone therapy, and therefore could profit from rescue-of- secretion therapy by ERQC/ERAD modulation, as a broad-spectrum alternative to pharmacological chaperone therapy. The work described here constitutes a proof-of-principle study, encouraging testing of modulators of ERQC, and adding to the ERAD modulation approach described in [12], towards the development of broad-spectrum rescue-of-secretion drugs for the therapy of congenital rare diseases associated with responsive glycoprotein mutants.

## Material and Methods

### Computational Methods

For the Molecular Dynamics (MD) simulations, we used with minor modifications the protocol described by Modenutti et al [59]. We utilised as initial structure a model built from the crystal structures of the cis-dimer of the extracellular domain of human Trop-2 (PDB IDs 7PEE [44] and 7E5N [45]) including only eight residues from the transmembrane region.

Starting from each structure (wild type and mutant), we prepared the systems and performed three replicas of a 1 µs all-atom MD simulation, using the AMBER package [60]. For each system, all non-protein molecules (carbohydrates) were removed from the PDB-deposited structure. The systems were prepared with the `tleap` module from the AMBER package, using the ff19SB/OPC force fields for amino acid/water molecules, respectively [61]. Standard protonation states were assigned to titratable residues (Asp and Glu are negatively charged; Lys and Arg are positively charged). Histidine protonation was assigned by favouring for each residue the formation of hydrogen bonds. All disulfide bonds as reported by Sun et al [45] were enforced/preserved throughout. The complete protonated systems were then solvated by a truncated cubic box of OPC waters, ensuring that the distance between the biomolecule surface and the box limit was at least 10 Å.

Each system was first optimised using a conjugate gradient algorithm for 5,000 steps, followed by 150 ps-long constant volume MD equilibration, in which the first 100 ps were used to gradually raise the temperature of the system from 0 to 300 K. The heating was followed by a 300 ps-long constant temperature and constant pressure MD simulation, to equilibrate the system density. During these temperature- and density-equilibration processes, the protein C_α_ atoms were constrained by a 5 kcal/mol/Å force constant, using a harmonic potential centred at each atom’s starting position. Next, a second equilibration MD of 500 ps was performed, in which the integration step was increased to 2 fs and the force constant for restrained C_α_s was decreased to 2 kcal/mol/Å, followed by 10 ns-long MD simulation with no constraints. Finally, a 1µs-long MD simulation was carried out, with no constraints and the ’Hydrogen Mass Repartition’ method, which allows an integration step of 4 fs [62].

All simulations were performed using the `pmemd.cuda` algorithm from the AMBER package [63]. Pressure and temperature were kept constant using the Monte-Carlo barostat and Langevin thermostats, respectively, using default coupling parameters. All simulations were performed with a 10 Å cutoff for nonbonded interactions, and periodic boundary conditions, using the Particle Mesh Ewald summation method for long-range electrostatic interactions. The `SHAKE` algorithm was applied to all hydrogen-containing bonds in all simulations with an integration step equal or higher than 2 fs. All trajectory processing and parameters calculations were performed with the `CPPTRAJ` [64] module of the AMBER package. Images of the molecules were prepared using the Visual Molecular Dynamics (VMD) program [65]. The plots were created using the `Matplotlib` Python library [66].

### Generation of *UGGT1*^-/-^, *UGGT2*^-/-^ and *UGGT1/2^-/-^* knock-out cell lines

HEK 293T and Vero E6 KO cell lines were generated as follows.

#### HEK 293T cells

*UGGT1*^-/-^, *UGGT2*^-/-^ and *UGGT1/2^-/-^*HEK 293T cells were generated using the CRISPR/Cas9 system from Addgene (http://www.addgene.org/). The sequences for guide RNAs were obtained from http://tools.genome-engineering.org and https://www.addgene.org/pooled-library/zhang-human-gecko-v2/. Guides (UGGT1: 5’- *gctgatgaacctccaccaga*-3’; UGGT2: 5’-*gacggtcgccgcgtccaagt*-3’) were synthesised by Microsynth AG (Switzerland). They were inserted into the lentiCRISPRv2-puro vector (Addgene 52961) using the BsmBI restriction site. The plasmids were transfected with Jet Prime (Polyplus) into HEK 293T cells cultured in DMEM supplemented with 10% FBS. For the generation of *UGGT1/2^-/-^* double KO cells, *UGGT1*-KO cells were re-transfected with the lentiCRISPRv2-puro containing the *UGGT2* guide and pCDNA3 vector to confer geneticin resistance. Two days after transfection, the cell culture medium was supplemented with 2

μg/mL puromycin for selection of the *UGGT1*^-/-^ and *UGGT2*^-/-^ clones or with 2% geneticin for selection of *UGGT1/2^-/-^* cells. Puromycin or geneticin-resistant clones were collected after 10 days and cultured in DMEM supplemented with 10% FBS. Lack of protein expression was confirmed by Western blotting.

#### Vero E6 cells

For the generation of the *UGGT1* and *UGGT2* KO Vero E6 cells (ATCC) we used CRISPR/Cas9 system as described in [67]. *Chlorocebus sabaeus* UGGT1 primers (Fwd1 CACCGGTGGCGTGAGGAGGCCTC Rev1 AAACTGAGGCCTCCTCACGCCACC; Fwd2 CACCGTGGCTGTTCTCCTCAGTAA Rev2 AAACTTACTGAGGAGAACAGCCAC) and *Cs* UGGT2 primers (Fwd1 CACCGCTGAAACATCTGAATAGC Rev1 AAACAGCTATTCAGATGTTTCAGC; Fwd2 CACCGTGATTCTCTATGCCGAATT Rev2 AAACAATTCGGCATAGAGAATCAC) were designed based on the *C. sabaeus* genome and cloned into pSpCas9(BB)-2A-Puro (a gift from Feng Zhang, Addgene # 62988) following the protocol in [68]. Vero E6 cells were transfected with the pair of CRISPR plasmids at 40% confluency using the Lipofectamine 3000 system (Thermo Fisher Scientific). Puromycin selection was performed between days 3-6 of culture at 33 μg/mL. The culture was separated into single cells at day 10 and clones propagated. Knockouts were confirmed by PCR screening, western blotting and sequencing (data not shown).

### Cell culture conditions

All of the applied cell lines were cultured in DMEM, high glucose, GlutaMAX™ media supplemented with 10% FBS and 1% Antibiotic-Antimycotic (AB/AM) (Thermo Fischer Scientific, Waltham, MA, USA). The cells were kept at 37°C in humidified atmosphere containing 5% CO_2_.

### Cloning of the WT and Q118E mutant Trop-2 into the pEYFP-N1 mammalian expression vector

The generation of the recombinant Trop-2-EYFP fusion constructs was carried out as described in [69]. The human WT Trop-2 and the Trop-2-Q118E mutant cDNAs [70] were amplified by PCR method (primers: F’ gcgatt**ctcgag**tccggtccgcgttcc-**XhoI** and R’ gcgcc**ggtacc**aagctcggttcctttc-**KpnI**) and sub-cloned into the pEYFP-N1 vector (Clontech, OH, USA) for mammalian expression to give the Trop-2-pEYFP-N1 and Trop-2-Q118E-pEYFP-N1 plasmids.

### UGGT1 re-complementation plasmids

Human UGGT1-encoding cDNA was cloned into the pCDNA3.1/Zeo(+) mammalian expression vector by using NheI/AflII restriction enzyme cloning sites. Additional 5’ (GCTAGCA) and ‘3 sequences (CTTAAG) were added to the cDNA to carry out the cloning procedure (GenScript, Piscataway, NJ, USA). The inactive UGGT1 D1454A variant was generated with direct mutagenesis by GenScript. The sequences of the constructs are provided in the supplementary documentation.

### Generation of the *UGGT1* and *UGGT1 D1454A* HEK 293T stable cell lines

For the generation of our stable cell lines, WT and *UGGT1^-/-^*HEK 293T cells were cultured in 6-well plates (Corning, NY, USA) with 500,000 cells/1.5 mL starting cell density in DMEM, high glucose, GlutaMAX™ media supplemented with 10% FBS without antibiotics. After 24 hours, the HEK 293T cells were transfected with 1 µg UGGT1-pCDNA3.1/Zeo(+) or UGGT1-D1454A-pCDNA3.1/Zeo(+) construct using X-tremeGENE 9 DNA transfection reagent (Roche, Basel, Switzerland) following the manufacturer’s instructions.

Stable transfectants were obtained by one-week selection with 400 µg/mL Zeocin (InvivoGen Europe, France), followed by one-week selection with 200 µg/mL Zeocin. After three weeks of selection, the cells were plated for the re-complementation experiments.

### Transient transfection with Trop-2-pEYFP-N1 and Trop-2-Q118E-pEYFP-N1 constructs

HEK 293T cells were cultured in 6-well plates (Corning, NY, USA) with 0.8x10^6^ cells/1.5 mL starting cell density in Gibco™ DMEM, high glucose, GlutaMAX™ media supplemented with 10% Gibco™ Fetal Bovine Serum (FBS) without antibiotics (Thermo Fischer Scientific, MA, USA). After 24 hours, the HEK 293T WT, *UGGT1*^-/-^, *UGGT2*^-/-^ and *UGGT1/2^-/-^*cells were transfected with 1 µg Trop-2-pEYFP-N1 or Trop-2-Q118E-pEYFP-N1 plasmids using X-tremeGENE 9 DNA transfection reagent (Roche, Basel, Switzerland) following the manufacturer’s instructions.

Vero E6 cells were cultured in 6-well plates (Corning, NY, USA) using Gibco™ DMEM, high glucose, GlutaMAX™ media supplemented with 10% Gibco™ Fetal Bovine Serum (FBS) without antibiotics. After 24 hours, the Vero E6 WT, *UGGT1*^-/-^, *UGGT2*^-/-^ and *UGGT1/2^-/-^* cells were transfected with 1 µg Trop-2-pEYFP-N1 or Trop-2-Q118E-pEYFP-N1 plasmids using Lipofectamine 3000 DNA transfection reagent following the manufacturer’s instructions (Life Technologies). Mock-transfection using the pEYFP-N1 plasmid was used as a negative control.

### Confocal microscopy analysis of the Trop-2-transfected HEK 293T and Vero E6 cell lines

For the confocal microscopy measurements, we used organelle-specific staining in live cells to visualise the cellular localisation of the EYFP-labelled Trop-2 proteins. All of the stock solutions were diluted in prewarmed phenol-red free DMEM high glucose media to the final concentration. ER-Tracker Red (BODIPY TR Glibenclamide) dye was used at 1 µM final concentration to mark the endoplasmic reticulum (data not shown). To visualise the plasma membrane, we applied Wheat Germ Agglutinin, Alexa Fluor 555 Conjugate (Thermo Fisher Scientific, Waltham, MA, USA) at 5 µg/mL working concentration. After 15 mins incubation at 37 °C, the supernatant was carefully replenished with prewarmed phenol red free, high glucose DMEM supplemented with 10% FBS. The nuclei were stained with NucBlue Live ReadyProbes Reagent (Thermo Fisher Scientific, Waltham, MA, USA) following the manufacturer’s instructions. After staining, we performed live cell imaging using a Zeiss LSM980 Airyscan 2 microscope equipped with a plan-apochromat 63x/1.4 objective, located at the Leicester University College of Life Sciences Advanced Imaging Facility (AIF). For live imaging, cells were incubated at 37°C and under 5% CO_2_. Images were acquired in confocal mode (pixel size 66 x 66 nm) using sequential channel settings to minimise bleed-through. All confocal images were taken with the same settings and single optical sections were analysed. Alternatively, after organelle staining, cells were fixed using 3.7% paraformaldehyde for 10 mins at 37°C. Quenching was performed using 50mM NH_4_Cl for 10 mins at RT. Nuclei were counterstained with DAPI and cellular localisation of EYFP-labelled Trop-2 proteins was imaged using a Zeiss LSM800 Airyscan 2 microscope equipped with a plan-apochromat 63x/1.4 objective, located at the University of Chieti.

### Analysis of the confocal microscopy images

For image analysis, we used Zeiss ZEN 3.1 Blue edition (Zeiss, Jena, Germany) and Fiji (version: 2.0.0-rc-69/1.52b) image processing pieces of software [46]. Zeiss LSM980 Airyscan 2 system with the Zen Blue software allows us to minimise bleed-through from the multiple channels by using sequential imaging and better emission wavelength separation. All of the confocal images were taken with the same settings and single optical sections were analysed. To measure the Trop-2 protein colocalization with the membrane dye, we performed multichannel intensity plot analysis by using Fiji image processing tools. A single line was drawn across the cells and the intensity profiles were measured by an intensity line plot tool macro for each channel (FILM Macro Toolsets, Imperial College London). The overlapping signals show the colocalization of different channels.

### Isolation of hydrophobic, membrane-associated proteins from HEK 293T cells with CelLytic^TM^ MEM Protein Extraction Kit

Twenty-four hrs post-transfection, the cells transfected with Trop-2-pEYFP-N1 and Trop-2-Q118E-pEYFP-N1 were harvested with 100 µL PBS containing 1mM EDTA and the cell suspensions were centrifuged down with 200 x g for 5 minutes. The separation of the hydrophobic and hydrophilic proteins from the pellet was carried out with CelLytic^TM^ MEM Protein Extraction Kit (Sigma-Aldrich, MO, USA) following the manufacturer’s instructions.

### Transient transfection of WT and *UGGT1*^-/-^ KO HEK 293F cell lines with Trop-2 expressing constructs

HEK 293F cells were cultured in FreeStyle™ 293 Expression Medium (Thermo Fisher Scientific, MA, USA). The initial concentration of the culture was 0.5x10^6^ cells/mL in 50 mL media. Cells were kept in an orbital shaker incubator at 37 °C, 120 rpm and 5% CO_2._ Twenty-four hrs later or when they reached the 1x10^6^ cells/mL concentration, the cells were transfected with 1 µg plasmid DNA per million cells using polyethylenimine (PEI) in 1:4 ratio respectively in the presence of 5 µM kifunensine. The DNA was diluted in 5 mL sterile PBS then the PEI was added to the PBS/DNA solution. After a 20 min incubation at room temperature, the cells were transfected with the mixture. The membrane-associated proteins were isolated with styrene-maleic acid (SMA) co-polymer 72-96 hrs post-transfection.

### Detergent-free isolation of membrane-associated proteins using styrene-maleic acid co-polymer lipid particles (SMALPs)

The cells were harvested 72-96 hrs post-transfection at 3,000 x g for 5 min. Lipodisq^®^ Styrene:Maleic Anhydride Copolymer 2:1 (Kit (Sigma-Aldrich, MO, USA) was used for the detergent-free membrane fraction purification. The applied protocol was based on [71]. Before using the pre-hydrated SMA co-polymer, we had to hydrate the SMA co-polymer, as described in [71]. The cellular pellet was diluted to 30 mg/mL in a modified Buffer 1 solution (20 mM Tris–HCl pH 8, 150 mM NaCl, 5 µM Tris(2-carboxyethyl)phosphine hydrochloride (TCEP), 10 % (v/v) glycerol) then 1% of hydrated co-polymer was added to the suspension. After 2-2.5 hrs of incubation at room temperature in an orbital shaker, the suspension becomes more transparent. The solubilized membrane proteins were separated from the cellular debris with ultracentrifugation at 100,000 x g for 30 minsat 4 °C.

### Western blot analysis of Trop-2 protein in various membrane fractions

Protein concentration was determined with Coomassie (Bradford) Protein Assay Kit (Thermo Fisher Scientific, MA, USA) to ensure that the same amount of protein sample was loaded in each lane. For the SDS-page, 4-12% NuPAGE^TM^ Bis-Tris precast polyacrylamide protein gels were used with 1x NuPAGE^TM^ MES SDS Running Buffer (Thermo Fisher Scientific, MA, USA). The samples were prepared with NuPAGE^TM^ LDS Sample Buffer (4X) (Thermo Fisher Scientific, MA, USA) in the presence or absence of a reducing agent (NuPAGE^TM^ Sample Reducing Agent (10X), Thermo Fisher Scientific, MA, USA) and denatured at 92 °C for 10 minutes. The proteins were transferred overnight onto methanol activated polyvinylidene difluoride membrane (PVDF, Sigma-Aldrich, MO, USA) in Transfer Buffer (25 mM Tris, 190 mM Glycine, 20% (v/v) methanol). After the transfer, the membrane was blocked with 1x TBST + 5% skimmed milk powder for 2 hrs at room temperature. The blocking was followed by multiple washing cycles with 1x TBST. GFP Monoclonal (GF28R) primary antibody (Thermo Fisher Scientific, MA, USA) was applied with 1:3000 dilution factor in 1x TBST+5% skimmed milk powder for 1.5 hrs at room temperature. After the 1x TBST washing cycles, Goat anti-Mouse IgG (H+L) Secondary Antibody DyLight 800 4x PEG (Thermo Fisher Scientific, MA, USA) was applied with 1:10,000 dilution factor in 1x TBST + 5% skimmed milk powder for 1 hr at room temperature. The membrane was washed after the secondary antibody and LI-COR Odyssey device was used with Image Studio^TM^ software (LI-COR Biosciences, NE, USA) for the detection.

### Transfection of Trop-2-EYFP and Q118E mutant into *ALG6^-/-^* background cells

The generation of the *ALG6^-/-^, ALG6/UGGT1^-/-^, ALG6/UGGT2^-/-^ ALG6/UGGT1/2^-/-^* cells has been previously described [20]. For each cell line 1.5 million cells were plated onto 6 cm plates and grown in DMEM (Sigma D5796) containing 10% FBS for 24 hours. The following day the cells were transfected with either the Trop-2-pEYFP-N1 or Trop-2-Q118E-pEYFP-N1 plasmids. Briefly, 4.8 μg of DNA was diluted in Opti-Mem^TM^ (Gibco 31985070) to a total volume of 80 μL. In a separate tube 12 μL of PEI (Polyscience 24765), formulated at 1 mg/mL, was added to 68 μL of Opti-Mem^TM^ and incubated at room temperature for 5 minutes. After incubation, the PEI containing solution was added to the diluted DNA and incubated for an additional 20 minsat 25 °C. The entire 160 μL of PEI:DNA mixture was added drop wise to each plate. The cells were returned to the incubator and incubated for 24 hrs at 37 °C and 5% CO_2_.

### Cellular glucosylation assay

Prior to harvesting the media of transfected cells was removed and 1 mL of fresh media containing 500 μM DNJ (Cayman 21065) was added. The cells were incubated for 5 hrs at 37 °C and 5% CO_2_. Before lysis, the media was removed, and the cells were washed with 1 mL of cold PBS and placed on ice. 500 μL of lysis buffer (20 mM MES, 100 mM NaCl, 30 mM Tris pH 7.5, 0.5% Triton X-100) containing both protease inhibitor (Thermo 1861278) and 20 mM N-ethylmaleimide was added to each plate and scraped to remove the attached cells. The lysate was removed and shaken vigorously for 10 min at 4 °C before centrifuging for 10 min at 20,817 x g and 4 °C. The centrifuged lysate was transferred to a fresh tube and normalised to 750 μL. To each normalised tube, 25 μL of glutathione resin (Cytiva 17075601) was added, and tubes were incubated for 1 hrs at 4 °C under gentle rotation to pre-clear the samples.

During this time glutathione resin pre-incubated with either GST-CRT or GST-CRT(Y109A) was prepared [20]. For each sample 50 μL of glutathione resin was equilibrated with lysis buffer. After equilibration the resin was split into two and 50 μg of recombinant GST-CRT or GST-CRT(Y109A) was added. The mixture was brought up to 500 μL with lysis buffer before incubating for 3 hrs under gentle agitation at 4 °C. After incubation, the beads were centrifuged at 950 x g at 4 °C for 5 min and washed twice with lysis buffer. The pre-cleared supernatant was also spun at 950 x g at 4 °C for 5 min to pellet the resin. The pre-cleared sample was then divided: 150 μL (20%) into 600 μL of cold acetone for the whole cell lysate and 262.5 μL (35%) to the GST-CRT and GST-CRT(Y109A) tubes. 200 μL of lysis buffer was added to each CRT and CRT(Y109A) tube and left to mix gently overnight at 4 °C. The tubes containing the whole cell lysate were mixed vigorously and incubated at -20°C overnight. The next day the glutathione resin was spun at 950 x g for 5 min at 4 °C and washed twice with 500 μl of lysis buffer without protease inhibitor. The acetone precipitated samples were spun for 10 min at 20,817 x g and 4 °C to pellet the precipitated protein. The acetone was removed, and the samples were dried for 5 min at 65 °C. 40 μL of loading buffer (30 mM Tris-HCl pH 6.8, 9% SDS, 15% glycerol, 0.05% bromophenol blue) containing 100 mM dithiothreitol (Sigma D9779) was added to each tube and the samples were boiled for 10 mins. 15 μl of each sample was resolved by 9% SDS-PAGE and transferred to a PVDF membrane (Millipore IPFL00010) and blotted using an anti-EYFP antibody (Thermo MA5- 15256) and an anti-mouse secondary antibody (LI-CORE 926-32210).

ImageJ was used to quantify the immunoblots. To determine the percentage of glucosylation the amount of protein found in the whole cell lysate (WCL), GST-CRT, and GST-CRT(Y109A) was first normalised by their input percentages. Any background from the GST-CRT(Y109A) was subtracted from the GST-CRT before being divided by the normalised protein found within the WCL and multiplied by 100. The resulting values represented the amount of WT or mutant Trop-2 glucosylated by UGGT.

### Flow cytometry analysis of membrane-recruited WT and Q118E Trop-2 mutants

*UGGT1^-/-^* Vero E6 cells transfected with Trop-2-pEYFP-N1 and Trop-2-Q118E- pEYFP-N1 plasmids were stained after 48 hrs for flow cytometry. An anti Trop-2 mAb (T16) and and anti Trop-2 pAb (AF650 from R&D Systems) were used to perform staining of live, non-permeabilized cells. The secondary antibodies were APC-labelled Donkey anti mouse (for binding to T16) or Donkey anti Goat (for binding to pAb) and were utilised for secondary staining of the cells after removal of the primary antibodies with 3 washes using PBS. Fluorescent signals in the EYFP and in the APC channels were acquired for subsequent evaluation of the MFIs.

## Author contributions

P.R. and M.T. conceived the study. G.T. and M.T. carried out cloning of the vectors for transfections established the transfection and fluorescence microscopy protocols. G.T., A.L., A.S., P.R. and C.J.H. carried out the biochemical characterisation of the WT and mutant glycoproteins. G.T. and K.R.S. carried out the fluorescence microscopy measurements on HEK 293T cells. G.T., K.R.S. and M.T. analysed the fluorescence microscopy data. K.P.G. and D.N.H. assayed reglucosylation *in cellula*. D.N.H., K.P.G., M.M. and T.S. generated the KO HEK 293T cells. J.C.H., S.V. and N.Z. generated the Vero E6 KO cells. M.T. carried out the fluorescence microscopy measurements on Vero E6 cells and the flow cytometry experiments. C.P.M. and P.R. carried out the *in silico* work. All authors were involved in the data analysis and the writing of the manuscript.

## Supplement

Gene name: *Hs*UGGT1 cDNA sequence

Length:4680bp

Additional 5’ sequence: GCTAGC

Additional 3’ sequence: CTTAAG

Sequence:

ATGGGCTGCAAGGGAGACGCGAGCGGTGCGTGTGCCGCGGGTGCGCTGCCGGTGACAGGAGTTTGCTATAAAATGGGAGTTCTGGTTGTACTCACTGTTCTGTGGCTGTTCTCCTCAGTAAAGGCCGACTCAAAAGCCATTACAACCTCTCTTACAACAAAATGGTTTTCCACTCCATTGTTGTTAGAAGCCAGTGAGTTTTTAGCAGAAGACAGTCAAGAGAAATTTTGGAATTTTGTAGAAGCCAGTCAAAATATTGGATCATCAGATCATGACGGTACCGATTATTCCTACTATCATGCAATATTGGAGGCTGCATTTCAGTTTCTGTCACCCCTCCAGCAGAATTTGTTTAAATTTTGTCTGTCCCTTCGTTCTTACTCAGCTACAATCCAAGCCTTCCAGCAGATAGCAGCTGATGAACCTCCACCAGAAGGATGTAATTCGTTTTTTTCAGTGCATGGAAAGAAGACTTGTGAATCTGATACCCTTGAGGCTCTTCTACTGACAGCCTCTGAAAGACCCAAACCTTTATTGTTCAAAGGAGATCACAGATATCCCTCGTCTAATCCTGAAAGCCCTGTGGTGATTTTCTACTCTGAGATTGGCTCTGAGGAATTTTCCAATTTTCACCGCCAGCTTATATCAAAAAGCAATGCAGGCAAAATCAATTATGTATTCAGACATTATATATTTAATCCCAGGAAGGAGCCTGTTTACCTCTCTGGCTATGGCGTGGAATTGGCCATTAAGAGCACTGAGTACAAGGCCAAGGATGATACTCAGGTGAAAGGAACTGAGGTAAACACCACAGTGATTGGTGAAAATGATCCTATTGATGAGGTTCAGGGGTTCCTCTTTGGAAAATTAAGAGATCTGCACCCCGACCTGGAGGGACAGTTGAAAGAACTCAGAAAGCATCTTGTAGAGAGCACCAATGAAATGGCACCTTTAAAGGTTTGGCAGTTGCAAGATCTCAGTTTCCAGACTGCTGCTCGAATCTTGGCTTCTCCTGTTGAGTTGGCTTTGGTTGTCATGAAGGATCTTAGTCAGAATTTTCCTACCAAAGCCAGAGCAATAACAAAAACAGCTGTGAGCTCAGAACTTAGAACCGAAGTGGAAGAGAATCAGAAGTATTTCAAGGGAACTTTAGGATTACAACCTGGAGATTCAGCCCTCTTCATCAATGGACTTCACATGGATTTAGATACACAGGATATATTCAGTCTGTTTGATGTGTTGAGGAATGAAGCTCGGGTAATGGAGGGTCTGCATAGATTGGGAATAGAAGGCCTTTCTCTGCATAATGTTTTGAAGCTGAACATCCAGCCCTCTGAGGCAGACTATGCCGTAGACATCCGGAGTCCTGCTATTTCATGGGTCAACAACCTGGAGGTTGATAGCAGATATAATTCGTGGCCTTCTAGTTTACAAGAGTTGCTTCGACCCACCTTTCCTGGTGTTATTCGGCAGATCAGGAAAAACTTACATAATATGGTTTTCATAGTTGATCCTGCACATGAGACCACAGCAGAGTTGATGAACACAGCTGAGATGTTCCTTAGTAATCATATACCACTAAGAATTGGTTTTATCTTTGTGGTTAATGACTCTGAAGATGTTGATGGGATGCAAGATGCTGGAGTGGCTGTTCTTAGAGCATATAATTATGTTGCCCAAGAAGTGGATGATTATCATGCCTTCCAGACTCTGACACATATCTATAACAAGGTGAGGACTGGAGAAAAAGTGAAAGTTGAACATGTGGTCAGTGTCCTGGAGAAGAAATATCCGTATGTAGAAGTGAATAGCATTTTGGGGATTGATTCTGCTTATGATCGGAATCGGAAGGAAGCAAGAGGCTACTATGAGCAGACTGGAGTTGGACCTCTGCCCGTTGTGCTGTTCAATGGAATGCCCTTTGAAAGGGAACAGCTAGACCCTGATGAGTTAGAAACCATCACAATGCATAAAATCCTGGAGACCACCACCTTCTTCCAAAGAGCGGTGTACTTGGGTGAACTGCCCCATGATCAAGATGTGGTAGAGTATATCATGAATCAGCCAAATGTTGTTCCACGAATCAATTCTAGGATTTTGACAGCTGAACGAGACTACCTGGATTTAACAGCGAGTAATAACTTCTTTGTGGATGATTATGCTAGATTTACTATCTTGGATTCCCAAGGCAAGACTGCTGCTGTAGCCAATAGTATGAACTATCTGACAAAGAAAGGAATGTCCTCCAAGGAAATCTATGATGATTCTTTTATTAGGCCAGTAACTTTTTGGATTGTTGGGGATTTTGATAGCCCTTCTGGACGGCAGTTACTGTATGATGCCATCAAACATCAGAAATCCAGTAACAATGTTAGAATAAGCATGATCAATAATCCTGCCAAAGAGATAAGCTATGAGAACACTCAGATCTCCAGAGCAATCTGGGCAGCTCTCCAAACTCAGACTTCCAACGCTGCTAAGAACTTCATCACCAAAATGGCCAAGGAGGGGGCTGCAGAGGCCCTGGCTGCAGGAGCTGACATTGCGGAGTTCTCTGTTGGGGGAATGGATTTCAGTCTTTTTAAAGAGGTCTTTGAGTCTTCCAAAATGGATTTCATTTTGTCTCATGCCGTGTACTGCAGGGATGTTCTGAAGCTGAAGAAGGGACAGAGGGCAGTGATCAGCAATGGAAGGATCATTGGGCCACTGGAGGATAGTGAGCTCTTTAATCAAGACGATTTCCACCTCCTCGAAAATATCATCTTAAAAACCTCAGGACAGAAAATAAAATCTCATATTCAACAGCTTCGGGTAGAAGAAGATGTGGCAAGCGACTTGGTAATGAAGGTGGATGCTCTTCTGTCAGCGCAACCAAAAGGAGATCCAAGAATCGAGTACCAGTTTTTTGAAGACAGACACAGTGCAATCAAACTGAGGCCGAAGGAAGGGGAGACATACTTTGATGTTGTGGCTGTCGTTGACCCTGTCACCAGAGAAGCACAGAGACTTGCTCCTTTGCTCTTGGTTTTGGCTCAGCTGATAAACATGAATCTGAGAGTATTTATGAACTGCCAATCCAAACTTTCTGACATGCCTTTAAAAAGCTTTTACCGTTATGTCTTAGAACCAGAGATTTCTTTCACTTCAGACAATAGTTTTGCTAAGGGTCCAATCGCAAAATTTTTGGATATGCCTCAGTCTCCACTGTTCACTCTGAATTTGAACACACCTGAGAGCTGGATGGTAGAATCTGTCAGAACACCATATGATCTTGATAATATTTATTTAGAAGAGGTGGACAGTGTAGTGGCTGCTGAGTATGAGCTGGAATACCTGTTACTGGAAGGTCATTGCTACGACATCACCACAGGCCAGCCTCCA

CGGGGACTACAGTTTACCTTAGGAACTTCAGCCAACCCGGTCATTGTGGACACCATTGTTATGGCCAATCTGGGCTACTTTCAGCTGAAAGCCAACCCAGGAGCTTGGATCCTCAGACTTAGGAAGGGACGCTCTGAAGATATTTATAGAATTTACAGCCACGATGGCACTGATTCTCCCCCTGATGCTGATGAGGTGGTTATCGTCCTCAACAACTTCAAAAGCAAAATTATTAAAGTGAAGGTTCAGAAGAAGGCAGATATGGTGAACGAAGACTTGCTGAGTGATGGAACGAGTGAGAATGAATCTGGATTTTGGGATTCCTTCAAATGGGGCTTTACAGGACAGAAGACTGAGGAAGTGAAGCAAGATAAAGATGACATAATTAATATTTTCTCCGTTGCATCTGGTCATCTCTACGAAAGATTTCTTCGCATAATGATGCTATCCGTGCTGAAGAATACCAAGACTCCTGTGAAATTCTGGTTCTTGAAGAATTACTTGTCCCCCACATTTAAGGAGTTTATACCTTACATGGCAAATGAATACAATTTCCAGTATGAGCTTGTTCAGTACAAATGGCCCCGGTGGCTTCATCAACAAACTGAAAAACAGCGTATCATCTGGGGTTACAAGATCCTCTTCCTGGATGTACTTTTCCCACTAGTTGTTGACAAGTTCCTGTTTGTGGATGCTGATCAGATTGTACGAACAGATCTGAAAGAGTTAAGAGATTTCAATTTGGATGGTGCTCCTTATGGTTACACTCCTTTCTGTGACAGCCGAAGAGAAATGGACGGCTACAGGTTCTGGAAGTCAGGGTACTGGGCCAGTCATTTAGCCGGGCGAAAGTATCATATCAGTGCACTATATGTTGTGGATCTGAAGAAGTTTAGGAAAATAGCTGCTGGTGACAGACTCAGGGGACAGTACCAAGGTCTGAGTCAGGACCCTAACAGCCTTTCAAATCTTGATCAAGATCTGCCCAATAACATGATTCATCAGGTGCCAATTAAATCCCTCCCTCAAGAATGGCTTTGGTGTGAAACGTGGTGTGATGACGCCTCTAAGAAAAGGGCAAAAACCATTGATTTGTGTAATAATCCGATGACCAAAGAGCCGAAACTGGAAGCAGCTGTGCGGATTGTCCCGGAGTGGCAGGACTACGACCAA GAGATCAAACAGCTACAGATCCGCTTTCAGAAGGAGAAAGAAACGGGAGCACTGTACAAAGAGAAGACA AAAGAACCAAGCCGAGAAGGTCCTCAGAAACGTGAAGAATTATGA

Cloning site: NheI/AflII

### The inactive mutants were generated with direct mutagenesis

*Hs*UGGT1 D1454A mutant_pcDNA3.1/Zeo (+)

Variant sequence:

GCTAGCATGGGCTGCAAGGGAGACGCGAGCGGTGCGTGTGCCGCGGGTGCGCTGCCGGTGACAGGAGTT TGCTATAAAATGGGAGTTCTGGTTGTACTCACTGTTCTGTGGCTGTTCTCCTCAGTAAAGGCCGACTCA AAAGCCATTACAACCTCTCTTACAACAAAATGGTTTTCCACTCCATTGTTGTTAGAAGCCAGTGAGTTT TTAGCAGAAGACAGTCAAGAGAAATTTTGGAATTTTGTAGAAGCCAGTCAAAATATTGGATCATCAGATCATGACGGTACCGATTATTCCTACTATCATGCAATATTGGAGGCTGCATTTCAGTTTCTGTCACCCCTCCAGCAGAATTTGTTTAAATTTTGTCTGTCCCTTCGTTCTTACTCAGCTACAATCCAAGCCTTCCAGCAGATAGCAGCTGATGAACCTCCACCAGAAGGATGTAATTCGTTTTTTTCAGTGCATGGAAAGAAGACTTGTGAATCTGATACCCTTGAGGCTCTTCTACTGACAGCCTCTGAAAGACCCAAACCTTTATTGTTCAAAGGAGATCACAGATATCCCTCGTCTAATCCTGAAAGCCCTGTGGTGATTTTCTACTCTGAGATTGGCTCTGAGGAATTTTCCAATTTTCACCGCCAGCTTATATCAAAAAGCAATGCAGGCAAAATCAATTATGTATTCAGACATTATATATTTAATCCCAGGAAGGAGCCTGTTTACCTCTCTGGCTATGGCGTGGAATTGGCCATTAAGAGCACTGAGTACAAGGCCAAGGATGATACTCAGGTGAAAGGAACTGAGGTAAACACCACAGTGATTGGTGAAAATGATCCTATTGATGAGGTTCAGGGGTTCCTCTTTGGAAAATTAAGAGATCTGCACCCCGACCTGGAGGGACAGTTGAAAGAACTCAGAAAGCATCTTGTAGAGAGCACCAATGAAATGGCACCTTTAAAGGTTTGGCAGTTGCAAGATCTCAGTTTCCAGACTGCTGCTCGAATCTTGGCTTCTCCTGTTGAGTTGGCTTTGGTTGTCATGAAGGATCTTAGTCAGAATTTTCCTACCAAAGCCAGAGCAATAACAAAAACAGCTGTGAGCTCAGAACTTAGAACCGAAGTGGAAGAGAATCAGAAGTATTTCAAGGGAACTTTAGGATTACAACCTGGAGATTCAGCCCTCTTCATCAATGGACTTCACATGGATTTAGATACACAGGATATATTCAGTCTGTTTGATGTGTTGAGGAATGAAGCTCGGGTAATGGAGGGTCTGCATAGATTGGGAATAGAAGGCCTTTCTCTGCATAATGTTTTGAAGCTGAACATCCAGCCCTCTGAGGCAGACTATGCCGTAGACATCCGGAGTCCTGCTATTTCATGGGTCAACAACCTGGAGGTTGATAGCAGATATAATTCGTGGCCTTCTAGTTTACAAGAGTTGCTTCGACCCACCTTTCCTGGTGTTATTCGGCAGATCAGGAAAAACTTACATAATATGGTTTTCATAGTTGATCCTGCACATGAGACCACAGCAGAGTTGATGAACACAGCTGAGATGTTCCTTAGTAATCATATACCACTAAGAATTGGTTTTATCTTTGTGGTTAATGACTCTGAAGATGTTGATGGGATGCAAGATGCTGGAGTGGCTGTTCTTAGAGCATATAATTATGTTGCCCAAGAAGTGGATGATTATCATGCCTTCCAGACTCTGACACATATCTATAACAAGGTGAGGACTGGAGAAAAAGTGAAAGTTGAACATGTGGTCAGTGTCCTGGAGAAGAAATATCCGTATGTAGAAGTGAATAGCATTTTGGGGATTGATTCTGCTTATGATCGGAATCGGAAGGAAGCAAGAGGCTACTATGAGCAGACTGGAGTTGGACCTCTGCCCGTTGTGCTGTTCAATGGAATGCCCTTTGAAAGGGAACAGCTAGACCCTGATGAGTTAGAAACCATCACAATGCATAAAATCCTGGAGACCACCACCTTCTTCCAAAGAGCGGTGTACTTGGGTGAACTGCCCCATGATCAAGATGTGGTAGAGTATATCATGAATCAGCCAAATGTTGTTCCACGAATCAATTCTAGGATTTTGACAGCTGAACGAGACTACCTGGATTTAACAGCGAGTAATAACTTCTTTGTGGATGATTATGCTAGATTTACTATCTTGGATTCCCAAGGCAAGACTGCTGCTGTAGCCAATAGTATGAACTATCTGACAAAGAAAGGAATGTCCTCCAAGGAAATCTATGATGATTCTTTT

ATTAGGCCAGTAACTTTTTGGATTGTTGGGGATTTTGATAGCCCTTCTGGACGGCAGTTACTGTATGATGCCATCAAACATCAGAAATCCAGTAACAATGTTAGAATAAGCATGATCAATAATCCTGCCAAAGAGATAAGCTATGAGAACACTCAGATCTCCAGAGCAATCTGGGCAGCTCTCCAAACTCAGACTTCCAACGCTGCTAAGAACTTCATCACCAAAATGGCCAAGGAGGGGGCTGCAGAGGCCCTGGCTGCAGGAGCTGACATTGCGGAGTTCTCTGTTGGGGGAATGGATTTCAGTCTTTTTAAAGAGGTCTTTGAGTCTTCCAAAATGGATTTCATTTTGTCTCATGCCGTGTACTGCAGGGATGTTCTGAAGCTGAAGAAGGGACAGAGGGCAGTGATCAGCAATGGAAGGATCATTGGGCCACTGGAGGATAGTGAGCTCTTTAATCAAGACGATTTCCACCTCCTCGAAAATATCATCTTAAAAACCTCAGGACAGAAAATAAAATCTCATATTCAACAGCTTCGGGTAGAAGAAGATGTGGCAAGCGACTTGGTAATGAAGGTGGATGCTCTTCTGTCAGCGCAACCAAAAGGAGATCCAAGAATCGAGTACCAGTTTTTTGAAGACAGACACAGTGCAATCAAACTGAGGCCGAAGGAAGGGGAGACATACTTTGATGTTGTGGCTGTCGTTGACCCTGTCACCAGAGAAGCACAGAGACTTGCTCCTTTGCTCTTGGTTTTGGCTCAGCTGATAAACATGAATCTGAGAGTATTTATGAACTGCCAATCCAAACTTTCTGACATGCCTTTAAAAAGCTTTTACCGTTATGTCTTAGAACCAGAGATTTCTTTCACTTCAGACAATAGTTTTGCTAAGGGTCCAATCGCAAAATTTTTGGATATGCCTCAGTCTCCACTGTTCACTCTGAATTTGAACACACCTGAGAGCTGGATGGTAGAATCTGTCAGAACACCATATGATCTTGATAATATTTATTTAGAAGAGGTGGACAGTGTAGTGGCTGCTGAGTATGAGCTGGAATACCTGTTACTGGAAGGTCATTGCTACGACATCACCACAGGCCAGCCTCCACGGGGACTACAGTTTACCTTAGGAACTTCAGCCAACCCGGTCATTGTGGACACCATTGTTATGGCCAATCTGGGCTACTTTCAGCTGAAAGCCAACCCAGGAGCTTGGATCCTCAGACTTAGGAAGGGACGCTCTGAAGATATTTATAGAATTTACAGCCACGATGGCACTGATTCTCCCCCTGATGCTGATGAGGTGGTTATCGTCCTCAACAACTTCAAAAGCAAAATTATTAAAGTGAAGGTTCAGAAGAAGGCAGATATGGTGAACGAAGACTTGCTGAGTGATGGAACGAGTGAGAATGAATCTGGATTTTGGGATTCCTTCAAATGGGGCTTTACAGGACAGAAGACTGAGGAAGTGAAGCAAGATAAAGATGACATAATTAATATTTTCTCCGTTGCATCTGGTCATCTCTACGAAAGATTTCTTCGCATAATGATGCTATCCGTGCTGAAGAATACCAAGACTCCTGTGAAATTCTGGTTCTTGAAGAATTACTTGTCCCCCACATTTAAGGAGTTTATACCTTACATGGCAAATGAATACAATTTCCAGTATGAGCTTGTTCAGTACAAATGGCCCCGGTGGCTTCATCAACAAACTGAAAAACAGCGTATCATCTGGGGTTACAAGATCCTCTTCCTGGATGTACTTTTCCCACTAGTTGTTGACAAGTTCCTGTTTGTGGATGCTGATCAGATTGTACGAACAGATCTGAAAGAGTTAAGAGATTTCAATTTGGATGGTGCTCCTTATGGTTACACTCCTTTCTGTGACAGCCGAAGAGAAATGGACGGCTACAGGTTCTGGAAGTCAGGGTACTGGGCCAGTCATTTAGCCGGGCGAAAGTATCATATCAGTGCACTATATGTTGTGGATCTGAAGAAGTTTAGGAAAATAGCTGCTGGTGACAGACTCAGGGGACAGTACCAAGGTCTGAGTCAGGACCCTAACAGCCTTTCAAATCTTGATCAAGCTCTGCCCAATAACATGATTCATCAGGTGCCAATTAAATCCCTCCCTCAAGAATGGCTTTGGTGTGAAACGTGGTGTGATGACGCCTCTAAGAAAAGGGCAAAAACCATTGATTTGTGTAATAATCCGATGACCAAAGAGCCGAAACTGGAAGCAGCTGTGCGGATTGTCCCGGAGTGGCAGGACTACGACCAAGAGATCAAACAGCTACAGATCCGCTTTCAGAAGGAGAAAGAAACGGGAGCACTGTACAAAGAGAAGACAAAAGAACCAAGCCGAGAAGGTCCTCAGAAACGTGAAGAATTATGACTTAAG

## Acknowledgments

P.R. was the recipient of a LISCB Wellcome Trust ISSF award, grant reference 204801/Z/16/Z. G.T. was funded by a Wellcome Trust Seed Award in Science to P.R., grant reference 214090/Z/18/Z. A.L. was the recipient of an Italian Government PhD Studentship. M.M. is supported by Signora Alessandra, AlphaONE Foundation, Foundation for Research on Neurodegenerative Diseases, Swiss National Science Foundation, Comel and Gelu Foundations. M.T. was supported by the Italian Ministry of University and Research, Programma Per Giovani Ricercatori “Rita Levi Montalcini” grant reference PGR12I7N1Z. We thank the Advanced Imaging Facility (RRID:SCR_020967) at the University of Leicester for support. The Zeiss LSM 980 Airyscan 2 system was funded by a BBSRC grant for the Advanced Imaging Facility (AIF), reference number BB/S019510/1. This work was also supported by the National Institutes of Health (GM086874 to D.N.H.) and a Chemistry-Biology Interface program training grant (T32 GM139789 to K.P.G.). Siyu Wang and Louise Fairall helped with HEK293F cell expression. Michelle Hill assisted us in the generation of the Vero E6 KO cell lines. N.Z. is a Fellow of Merton College Oxford.

---

**Figure S1.**
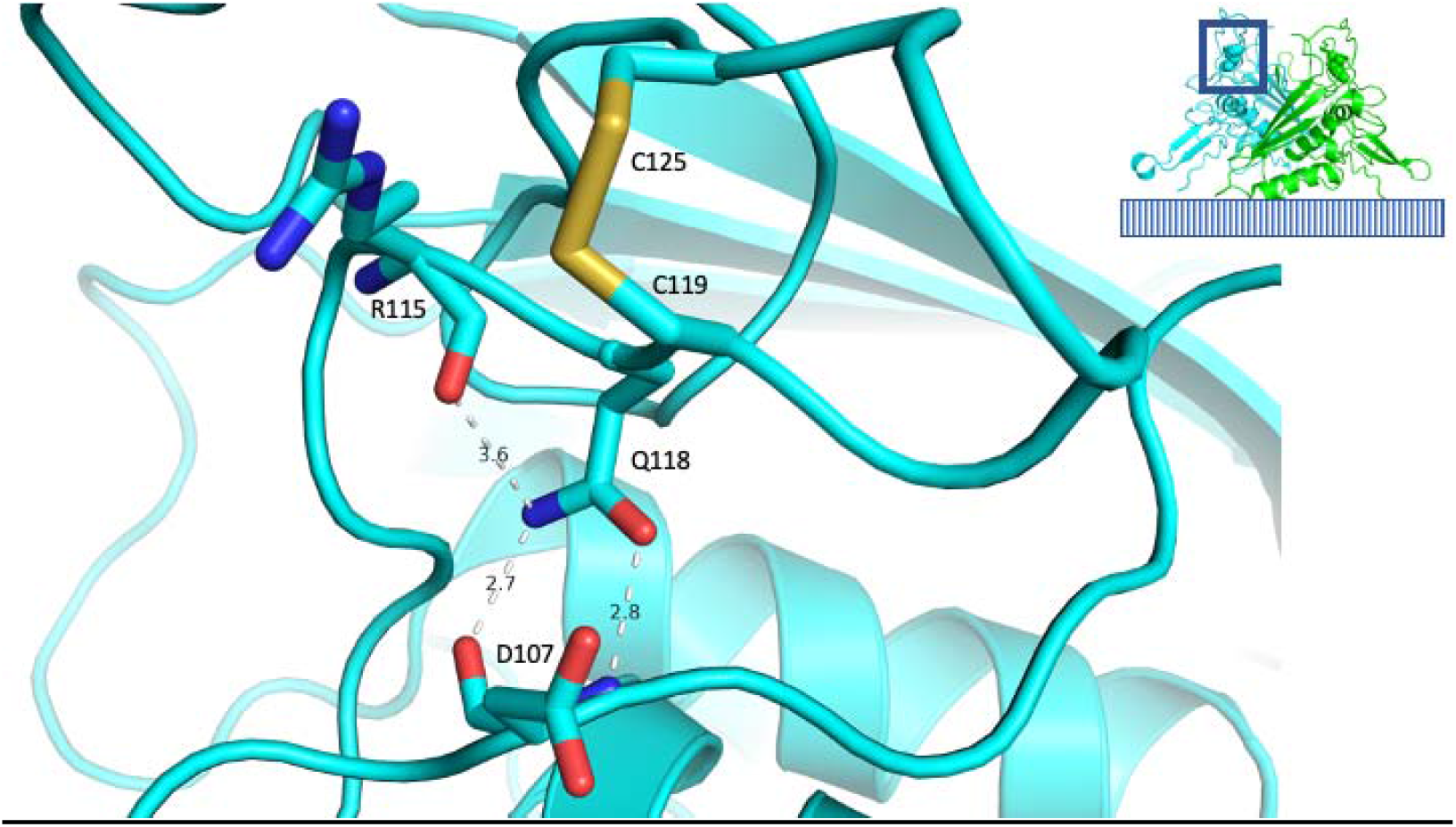
Native interactions of Q118 in the wild type structure of the Trop-2 dimer. The region of Trop-2 surrounding Q118 is shown in cyan cartoon representation. The Q118 residue and a few of its relevant neighbours are represented as sticks, coloured by atom type (carbon, oxygen and nitrogens in cyan, red and blue, respectively). A few interatomic distances between Q118 and neighbouring residues are indicated by white dashed lines and annotated with their length (in Å). The inset shows the Trop-2 extracellular domain dimer in cartoon representation, with the monomers coloured in cyan and green, Q118 in spheres representation, and a symbolic representation of the outer plasma membrane phospholipid layer on which the dimer sits. Based on PDB ID 7E5N. Figure made in PyMOL.

**Figure S2.**
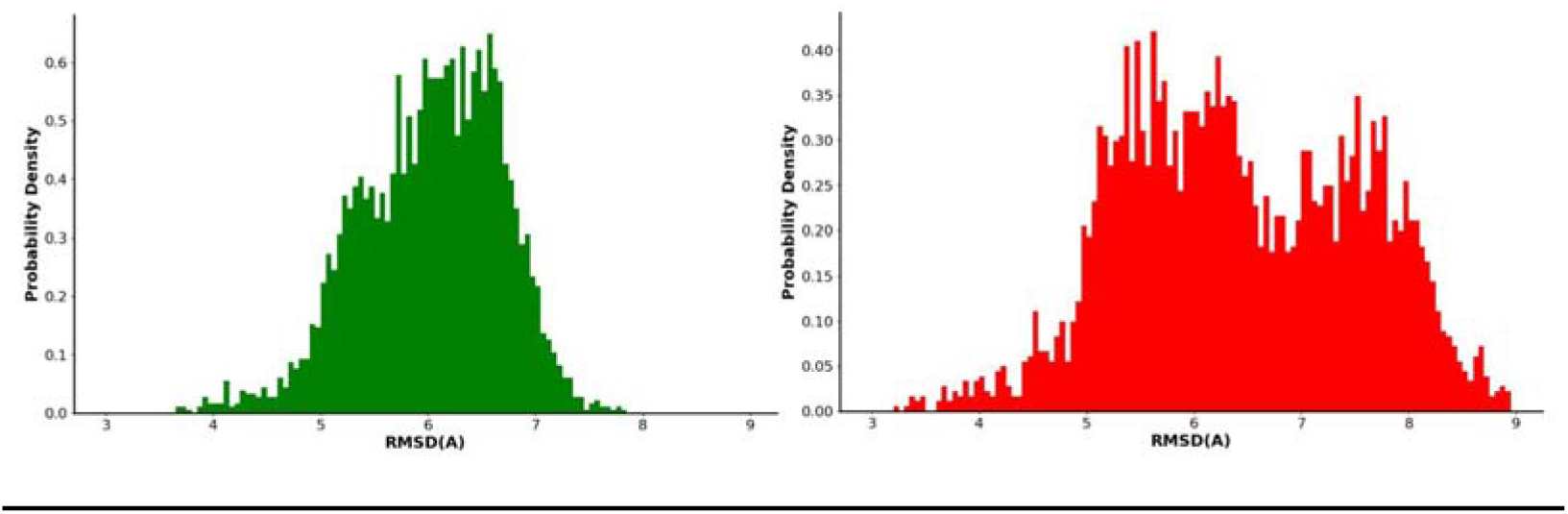
Average fluctuation of protein backbone rmsd_C__α_ during a 1 µs MD simulation. The histograms of the root mean square deviation from the initial model (rmsd_Cα_) of the Trop-2 extracellular domain dimer, as sampled during the MD simulations, are shown in green (WT) and red (Q118E mutant).

**Figure S3.**
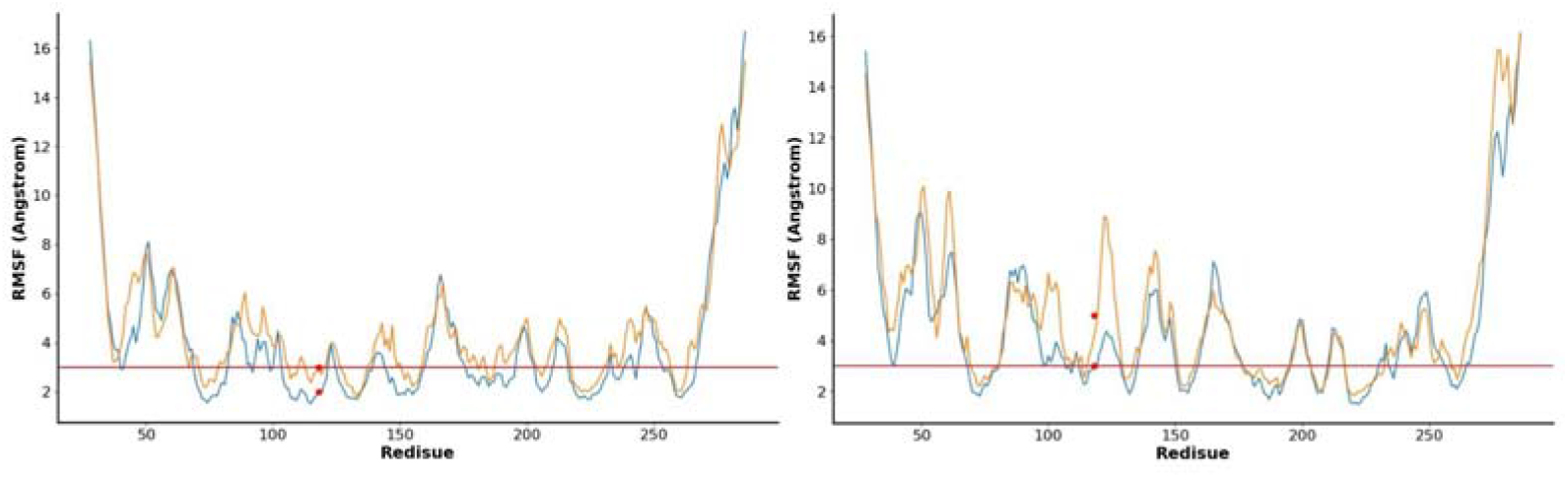
Per residue average fluctuation rmsf_C__α_ of protein backbone during a 1 µs MD simulation. The values for the root mean square fluctuation (rmsf_Cα_) for each position of the Trop-2 extracellular domain dimer, as sampled during the MD simulations, are shown as blue and orange lines for monomers one and two, respectively. The horizontal line indicates the average value for the wild type protein (3 Å, excluding C-ter and N-ter). The red dots indicate position 118.

**Figure S4.**
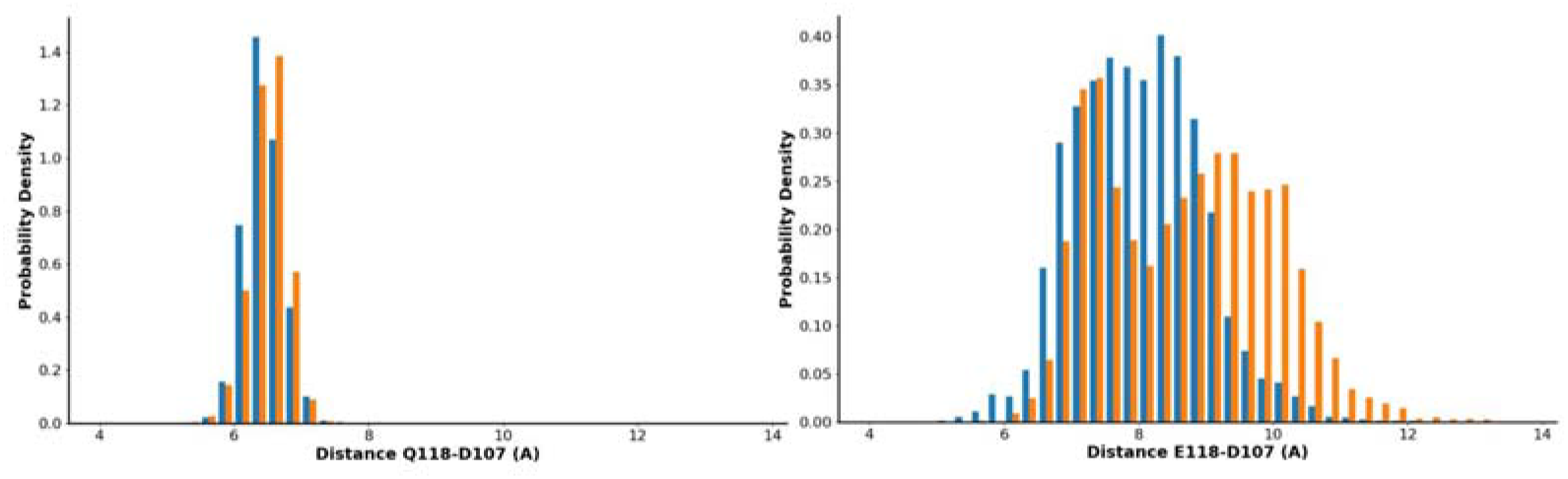
Effect of Q118E mutation over the local structure of the Trop-2 TY domain during a 1 µs MD simulation. The distance between the C_α_ residues of D107 and Q(E)118 are taken as representative of the local structure of the Trop-2 TY subdomain. The histograms of the values for the D107 C_α_ - Q(E)118 C_α_ distance, as sampled during the MD simulations, are shown in blue and orange for monomer one and two of the Trop-2 extracellular domain dimer, respectively.

## Notes

### Competing Interest Statement

The authors have declared no competing interest.

